# A Blood Immunological Signature of Subclinical Coronary Artery Atherosclerosis in People Living with HIV-1 Receiving Antiretroviral Therapy

**DOI:** 10.1101/2023.12.15.571922

**Authors:** Tomas Raul Wiche Salinas, Yuwei Zhang, Annie Gosselin, Natalia Fonseca Do Rosario, Mohamed El-Far, Ali Filali-Mouhim, Jean-Pierre Routy, Carl Chartrand-Lefebvre, Alan L. Landay, Madeleine Durand, Cécile L. Tremblay, Petronela Ancuta, the Canadian HIV and Aging Cohort Study (CHACS)

**Affiliations:** CHUM-Research Centre, Montréal, Qc, Canada; Département de microbiologie, infectiologie et immunologie, Faculté de médecine, Université de Montréal, Montréal, QC, Canada; Chronic Viral Illness Service and Division of Hematology, Research Institute of the McGill University Health Centre, Montreal, QC, Canada; Département de Radiologie, Radio-oncologie et Médecine Nucléaire, Faculté de Médecine, Université de Montréal, Montréal, QC, Canada; Rush University Medical Center, Chicago, IL, USA; Département de médecine, Faculté de médecine, Université de Montréal, Montréal, QC, Canada

**Keywords:** HIV-1, antiretroviral therapy (ART), cardiovascular disease (CVD), Th17/Treg cells, non-classical monocytes, myeloid/plasmacytoid dendritic cells

## Abstract

Cardiovascular disease (CVD) remains an important co-morbidity in people living with HIV-1 (PLWH) receiving antiretroviral therapy (ART). Our previous studies performed on the Canadian HIV/Aging Cohort Study (CHACS) (>40 years-old; Framingham Risk Score (FRS) >5%), revealed a 2-3-fold increase in non-calcified coronary artery atherosclerosis (CAA) plaque burden, measured by Computed tomography angiography scan (CTAScan) as total (TPV) and low attenuated plaque volume (LAPV) in ART-treated PLWH (HIV+) *versus* uninfected controls (HIV-). In an effort to identify novel correlates of subclinical CAA, markers of intestinal damage (sCD14, LBP, FABP2); cell trafficking/inflammation (CCL20, CX3CL1, MIF, CCL25); subsets of Th17-polarized and regulatory (Tregs) CD4^+^ T-cells, classical/intermediate/non-classical monocytes, and myeloid/plasmacytoid dendritic cells, were studied in relationship with HIV and TPV/LAPV status. The TPV detection/values coincided with higher plasma sCD14, FABP2, CCL20, MIF, CX3CL1 and triglyceride levels, lower Th17/Treg ratios, and classical monocyte expansion. Among HIV^+^, TPV^+^ *versus* TPV^−^ exhibited lower Th17 frequencies, reduced Th17/Treg ratios, higher frequencies of non-classical CCR9^low^HLADR^high^ monocyte, and increased plasma fibrinogen levels. Finally, Th17/Treg ratios and non-classical CCR9^low^HLADR^high^ monocyte frequencies remained associated with TPV/LAPV after adjusting for FRS and HIV/ART duration in a logistic regression model. These findings point to Th17 paucity and non-classical monocyte abundance as novel immunological correlates of subclinical CAA that may fuel the CVD risk in ART-treated PLWH.

## Introduction

Mortality in people living with human immunodeficiency virus type 1 (HIV) (PLWH) has considerably diminished after the implementation of antiretroviral therapy (ART). Nevertheless, non-AIDS co-morbidities, such cardiovascular disease (CVD), remain highly prevalent in ART-treated PLWH [1–8]. A modeling study performed in the Netherlands estimated an increase in the life expectancy of ART-treated PLWH from 43.9 years in 2010 to 56.6 years in 2030, and predicted that 78% of PLWH anticipated will be diagnosed with CVD [9,10]. Indeed, PLWH tend to present clinical signs of CVD approximately 10 years earlier compared to general population[11]. In addition to traditional CVD risk factors *(e.g.,* smoking, hypertension, dyslipidemia, diabetes, insulin resistance, and/or sedentary life-style) [12] and mental health disorders with an impact on CVD risk, HIV-specific mechanism (e.g., HIV-mediated metabolic alterations due to long-term administration of ART) contribute to the premature occurrence of CVD in ART-treated PLWH [7,8,10,13–15]. In addition, the persistence of HIV reservoirs during ART is associated with impaired intestinal mucosal barrier functions, microbial translocation, and immune dysfunction, and chronic inflammation, which, together with CMV and other co-infections, contribute to an increased CVD risk in this group [16–20]. Thus, CVD represents a major non-AIDS co-morbidity in ART-treated PLWH and novel therapeutic interventions are needed to reduce this risk.

Cells of the adaptive immune system are key players in CVD pathogenesis [12,21,22]. Among them, T lymphocytes infiltrate atherosclerotic plaques and are recruited into the heart *via* mechanisms involving the hepatocyte growth factor receptor c-Met, CCR4, CXCR3, and CCR5 [23]. The antigenic specificity of such CD4^+^ T-cell subsets remains yet unclear [24]. In HIV-uninfected individuals, T helper 1 (Th1) cells have pro-atherogenic features, whereas regulatory T-cells (Tregs) and Th17 cells were reported to play dual roles in the development of atherosclerosis, either protective or pathogenic [24,25]. In individuals with unstable angina or acute myocardial infarction, the frequency of Th17 cells was higher compared to those in participants with stable angina or healthy individuals [26]. The role of CD4^+^ T-cells and their associated cytokines in atherosclerosis or coronary artery disease in ART-treated PLWH remains poorly documented, with some recent studies documenting the expansion Tregs [27] and surprisingly Th17 cells [28]. Also, ART-treated PLWH with carotid atherosclerotic plaque, increased carotid intima media thickness, or arterial stiffness, presented with increased circulating CD8^+^ T-cell activation (CD38^+^HLA-DR) [29,30].

In addition to T cells, innate immune cells, such as monocytes and myeloid dendritic cell (mDC) also contribute to atherosclerotic plaque formation/rupture [31–33]. Mice deficient in CCR2 and CX3CR1, two chemokine receptors important for the migration of classical (CD14^++^CD16^−^) and intermediate (CD14^++^CD16^++^) and non-classical (CD14^dim^CD16^++^) monocyte, respectively, exhibit decreased atherosclerotic plaque severity thus, pointing to the deleterious role of these monocytes in CVD [34,35]. Studies in mice deficient in Nur77/NR4A1, that lack non-classical monocytes, reported controversial results that may be explained by differences in study design [35]. Nevertheless, human studies support a deleterious role of monocytes in CVD pathogenesis [35], as well as the pro-inflammatory features of non-classical monocytes [36,37]. In PLWH compared to uninfected individuals, arterial inflammation is higher and correlates with increased circulating levels of the pro-inflammatory cytokine IL-6 and monocytes activation [38,39]. In uninfected individuals, an elevated frequency of circulating non-classical monocytes was associated with an amplified carotid intima-media thickness (IMT) over 10 years in a prospective cohort study [40]. Similarly, in PLWH, the intermediate monocyte counts were associated to subclinical atherosclerosis [41] and the expression of CX3CR1 on CD16^+^ monocytes independently predicted carotid artery thickness [42]. Of note, a decreased expression of CXCR4 was observed on non-classical monocytes in women with subclinical atherosclerosis [43]. Moreover, *ex vivo* experiments with monocytes of virologically suppressed PLWH showed an increased potential to form atherosclerosis promoting foam cells compared to uninfected individuals [44]. Furthermore, atherosclerosis plaque burden was associated with increased levels of the monocyte chemoattractant protein-1 (MCP-1/CCL2) in ART-treated PLWH [45,46].

Other innate immune cells, such as plasmacytoid DC (pDC) are documented to infiltrate atherosclerotic lesions [47,48], but their contribution to atherosclerosis remains controversial. Some mice studies have shown that pDCs promote atherogenesis through their capacity to produce interferon (IFN)-α [49], while others support their role in atheroprotection [50]. pDCs contribute to peripheral and central tolerance role by the induction of Tregs; or, the anti-atherogenic role of pDCs was associated with indoleamine 2,3-dioxygenase 1 (IDO1)-dependent induction of aortic Treg cells [51]. During HIV infection, there is a decline in pDC counts and alterations in their functions, which are not restored with ART [52,53]. Whether interventions to restore pDC may limit the CVD risk in ART-treated PLWH requires investigations.

The objective of this study was to identify an immunological signature associated with subclinical coronary artery atherosclerosis (CAA) in ART-treated PLWH in local Canadian cohorts. To reach this objective, we had access to plasma and PBMC samples collected at baseline from ART-treated PLWH (HIV^+^; n=61) and HIV-uninfected participants (HIV^−^; n=21) included in the cardiovascular imaging sub-study of the Canadian HIV/Aging Cohort Study (CHACS), a multi-center prospective controlled cohort study initiated in 2011 [54,55]. Our results reveal alterations in Th17/Treg ratios and the frequencies of non-classical CCR9^low^HLA-DR^high^ monocytes, together with overt plasma fibrinogen levels, that all correlated with the magnitude of subclinical CAA in ART-treated PLWH. Of particular importance, the predictive CVD risk value of Th17/Treg ratios and non-classical CCR9^low^HLA-DR^high^ monocyte expansion remained valid upon adjustment to traditional CVD risk factors included in the Framingham risk score (FRS) (*i.e.,* smoking, statins, LDL, triglycerides), as well as HIV-specific parameters (HIV/ART duration). Our findings emphasize the need for novel therapies aimed at restoring Th17 paucity at mucosal level and reducing monocyte-mediated inflammation to limit the CVD risk in ART-treated PLWH.

## Methods

### Ethics statement

This study was approved by the Institutional Review Boards (IRB) of the Centre de recherche du Centre hospitalier de l’Université de Montréal (CRCHUM) (Ethical approval #CE.11.063) and the IRBs of all participating sites. This study was conducted with the ethical principles for medical research involving human subjects established by the Declaration of Helsinki in 1975. All study participants provided written informed consent for the collection of blood and the use of plasma and peripheral blood mononucleated cells (PBMC) for the current research investigation.

### Study Participants

The design of the Canadian HIV and Aging Cohort Study (CHACS) has been previously reported [54–56]. The first eligible HIV-(n=21) and HIV+ (n=61) participants recruited in the CHACS were included in this sub-study. Briefly, inclusion criteria for the cardiovascular imaging sub-study of the CHACS cohort were subjects older than 40 years old, without clinical manifestations or diagnosis of CAA at recruitment and a 10 year risk of cardiovascular disease according to the Framingham risk score (FRS, calculated based on age, HDL and cholesterol levels, systolic blood pressure, smoking, diabetes, statin treatment [54–56]) ranging from 5-20% (Table 1). Exclusion criteria were renal impairment and hypersensitivity to contrast agents. Plasma and PBMC samples collected at baseline from ART-treated PLWH (HIV^+^) and uninfected (HIV^−^) participants were available for this study.

**Table 1:**
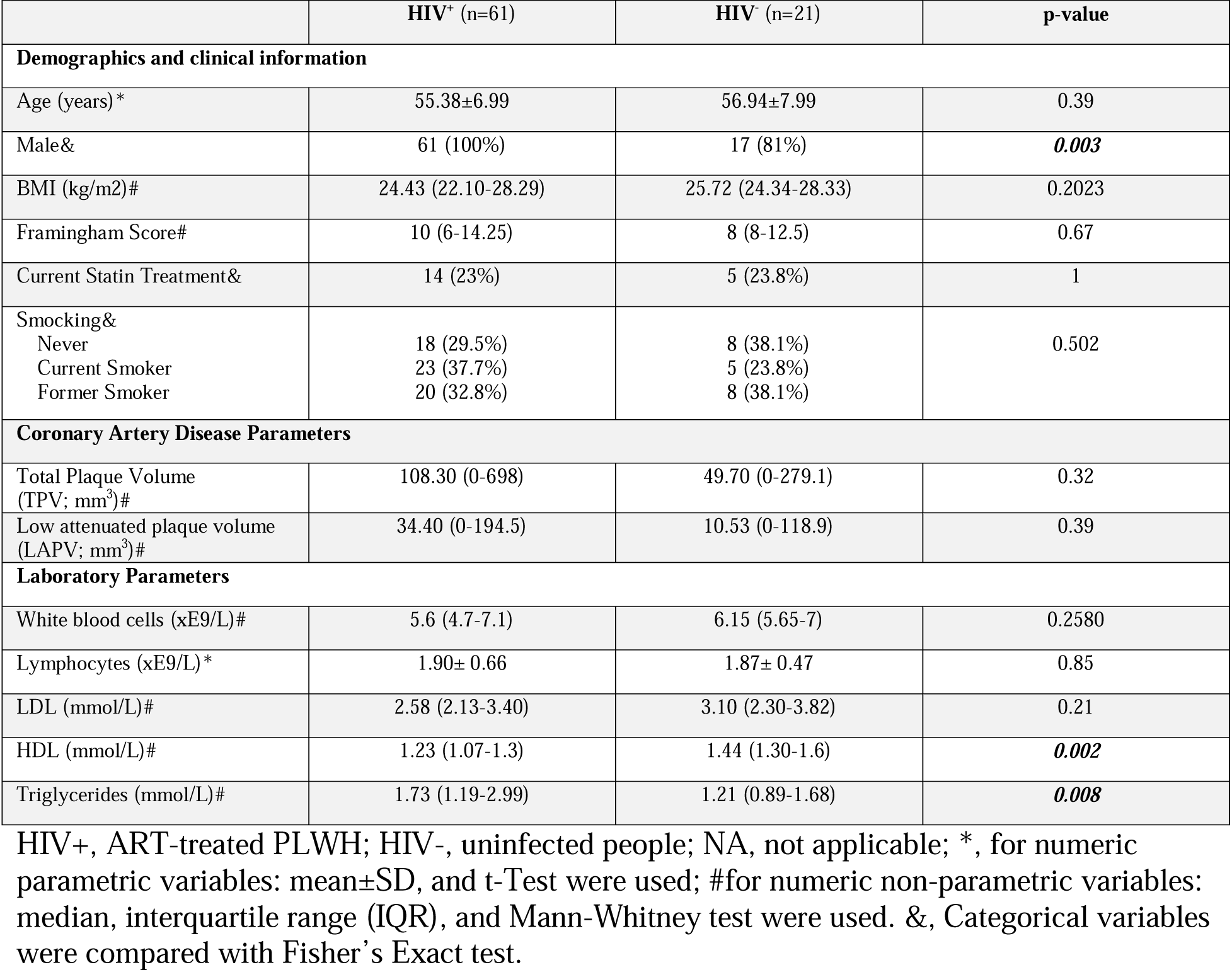
Description of study participants.

|  | HIV<sup>+</sup> (n=61) | HIV<sup>-</sup> (n=21) | p-value |
| --- | --- | --- | --- |
| **Demographics and clinical information** |  |  |  |
| Age (years)* | 55.38±6.99 | 56.94±7.99 | 0.39 |
| Male& | 61 (100%) | 17 (81%) | **0.003** |
| BMI (kg/m2)# | 24.43 (22.10-28.29) | 25.72 (24.34-28.33) | 0.2023 |
| Framingham Score# | 10 (6-14.25) | 8 (8-12.5) | 0.67 |
| Current Statin Treatment& | 14 (23%) | 5 (23.8%) | 1 |
| Smocking&<br>Never<br>Current Smoker<br>Former Smoker | 18 (29.5%)<br>23 (37.7%)<br>20 (32.8%) | 8 (38.1%)<br>5 (23.8%)<br>8 (38.1%) | 0.502 |
| **Coronary Artery Disease Parameters** |  |  |  |
| Total Plaque Volume (TPV; mm<sup>3</sup>)# | 108.30 (0-698) | 49.70 (0-279.1) | 0.32 |
| Low attenuated plaque volume (LAPV; mm<sup>3</sup>)# | 34.40 (0-194.5) | 10.53 (0-118.9) | 0.39 |
| **Laboratory Parameters** |  |  |  |
| White blood cells (xE9/L)# | 5.6 (4.7-7.1) | 6.15 (5.65-7) | 0.2580 |
| Lymphocytes (xE9/L)* | 1.90± 0.66 | 1.87± 0.47 | 0.85 |
| LDL (mmol/L)# | 2.58 (2.13-3.40) | 3.10 (2.30-3.82) | 0.21 |
| HDL (mmol/L)# | 1.23 (1.07-1.3) | 1.44 (1.30-1.6) | **0.002** |
| Triglycerides (mmol/L)# | 1.73 (1.19-2.99) | 1.21 (0.89-1.68) | **0.008** |

HIV<sup>+</sup>, ART-treated PLWH; HIV<sup>-</sup>, uninfected people; NA, not applicable; \*, for numeric parametric variables: mean±SD, and t-Test were used; #for numeric non-parametric variables: median, interquartile range (IQR), and Mann-Whitney test were used. &, Categorical variables were compared with Fisher's Exact test.

### Computed Tomography Angiography Scan

All study participants underwent Computed tomography angiography scan (CTA-Scan) to determine total plaque volume (TPV) and low attenuated plaque volume (LAPV, high-risk atherosclerotic plaque rupture), as we previously described [5,56–58]. Briefly, TPV represents the total volume (calculated from 3D reconstructions) of all coronary artery plaque present in an individual’s coronary arteries. Coronary arteries that are normal (free of signs of atherosclerosis) will have a TPV and LAPV of zero, while any volume greater than zero indicates the presence of CVD. For this sub-study, HIV^+^ and HIV^−^ participants were stratified based on TPV and LAPV values of zero or greater than zero, indicative of the absence or the presence of CVD, respectively (Table 2).

**Table 2:** Description of ART^−^treated PLWH with and without subclinical atherosclerosis.

|  | TPV<sup>+</sup> (n=39) | TPV<sup>-</sup> (n=22) | p-value |
| --- | --- | --- | --- |
| **Demographics and clinical parameters** |  |  |  |
| Age (years)* | 56.24±6.35 | 53.85±7.93 | 0.1625 |
| Male& | 39 (100%) | 22 (100%) | NA |
| BMI (kg/m<sup>2</sup>)# | 24.05 (21.05- 28.38) | 25.29 (22.91- 28.27) | 0.23 |
| Framingham Score# | 11.00 (7- 18) | 8.00 (5.5-13) | <u>0.085</u> |
| Current Statin Treatment& | 12 (30.8%) | 2 (9.1%) | <u>0.064</u> |
| Smocking&<br>Never<br>Current Smoker<br>Former Smoker | 7 (17.9%)<br>20 (51.3%)<br>12 (30.8%) | 11 (50%)<br>3 (13.6%)<br>8 (36.4%) | **0.005** |
| **Coronary Artery Disease Parameters** |  |  |  |
| Total Plaque Volume (mm<sup>3</sup>)# | 608.70 (142.6- 881.8) | 0 | NA |
| Low attenuated plaque volume (mm<sup>3</sup>)# | 164.80 (36- 296) | 0 | NA |
| **HIV disease parameters** |  |  |  |
| Undetectable Viral Load& | 37 (94.8%) | 19 (86.36%) | 0.34 |
| Duration of HIV (years)# | 19.16 (14.29- 24.4) | 14.71 (4.58- 24.14) | <u>0.06</u> |
| Duration of ART (years)* | 16.14±5.68 | 10.69±7.82 | **0.003** |
| Nadir CD4 (x10<sup>9</sup>/L)# | 195 (100- 330) | 170 (78- 255) | 0.397 |
| **Laboratory Parameters** |  |  |  |
| White blood cells (x10<sup>9</sup>/L)# | 5.90 (4.7- 7.8) | 5.40 (4.3- 6.85) | 0.27 |
| Lymphocytes (x10<sup>9</sup>/L)* | 1.91±0.60 | 1.89±0.78 | 0.924 |
| D-Dimer (ug/L)# | 300.00 (197.5- 410) | 290.00 (175- 380) | 0.52 |
| Fibrinogen (G/L)# | 3.23 (2.73- 3.78) | 2.67 (2.27- 3) | **0.005** |
| LDL (mmol/L)# | 2.40 (1.99- 3.48) | 2.90 (2.23- 3.40) | 0.35 |
| HDL (mmol/L)# | 1.21 (1.07- 1.37) | 1.25 (0.98- 1.37) | 0.96 |
| Triglycerides (mmol/L)# | 2.07 (1.26- 3.1) | 1.38 (1.09- 2.03) | 0.109 |
| **Serology** |  |  |  |
| Co-infection CMV# | 36 (92.3%) | 17 (77.3%) | 0.39 |

TPV, total plaque volume; NA, not applicable; \*, for numeric parametric variables: mean±SD, and t-Test were used; #, for numeric non-parametric variables: median, IQR, and Mann-Whitney test were used. &, Categorical variables were compared with Fisher's Exact test.

### ELISA

Markers of microbial translocation (LPS binding protein, LBP), mucosal damage (Intestinal-type Fatty Acid Biding Protein, I-FABP), immune activation (sCD14), and chemokines (CCL20, CX3CL1, CCL25, MIF) were quantified in plasma by ELISA, according with the manufacturer’s protocol (R&D Systems, Minneapolis, MN).

### Flow Cytometry

Fluorescence-activated cell sorting (FACS) was used to identify subsets of CD3^+^CD4^+^ T-cells [Th17 (CCR6^+^CD26^+^CD161^+^), Tregs (CD127^−^CD25^+^Foxp3^+^), central memory (CM, CD45RA^−^ CCR7^+^), effector memory (EM, CD45RA^−^CCR7^−^), effector memory RA^+^ (TEMRA, CD45RA^+^CCR7^−^), and naive (CD45RA^+^CCR7^+^) cells]; subsets of monocytes (CD3^−^CD4^low^HLA-DR^+^CD1c^−^) [classical (CD14^++^CD16^−^), intermediate (CD14^++^CD16^+^), non-classical (CD14^−^ CD16^++^), and Slan/M-DC8^+^]; as well as myeloid (mDC; HLA-DR^+^CD1c^+^) and plasmacytoid dendritic cells (pDC, BDCA2^+^CD123^+^); and measure their expression of chemokine receptors (CCR2, CCR6, CCR9, CX3CR1). Cells were analyzed by FACS using BD-LSRII cytometer, BD-Diva (BD Biosciences), and FlowJo version 10 (Tree Star, Inc., Ashland, Oregon, USA) software. The antibodies used in this study are presented in Supplemental Table 1. All Abs were titrated for an optimal noise/ratio. The gating strategy used to identify the different immune cell subsets is described in Supplemental Figures 1, 5 and 8. A combination of gates using the Boolean tool gating from Flow Jo was used to determine the expression of chemokines.

### Statistical Analysis

The Shapiro-Wilk test was used to determine the normal distribution of the continuous variables. Variables with normal distribution were presented as means with standard deviation, and variables with non-normal distribution were presented as median with interquartile range (IQR). Comparison of two groups for the same variable with normal distribution was analyzed using t-Test, and non-normal distributed variables were analyzed with Mann Whitney. Categorical variables were represented as proportions, and comparisons for the same variable between groups were analyzed using Fisher’s Exact Test. To assess the effect of potential confounding factors on the association between the immune subset and presence or absence of coronary plaque, logistic regression analyses were implemented in R (https://cran.r-project.org). First, we compared univariate and multivariate logistic regression models to study the contribution of the immune subset alone and in the presence of four sets of potential confounding factors (Models 1-4), as well as fibrinogen alone (Table 3). Model 1 adjusted for ART duration, smoking and the use of statins. Model 2 adjusted for ART duration, smoking and LDL. Model 3 adjusted for ART duration, smoking and triglycerides. Model 4 adjusted for ART duration, HIV duration and FRS. The confounding effect was estimated by the measure of the association (OR, odds ratio) before and after adjusting (% Δ (delta) OR). A change in the estimated association measure of less than 10% was used to exclude confounders, while a change superior to 30% identified large confounding bias [59]. P-values adjustment for multiple testing hypothesis was performed according to the method of Benjamin and Hochberg [60], which controls the false discovery rate with adjusted p-value cutoffs of 0.05.

**Table 3:** Logistic regression prediction analysis for the association between fibrinogen and cell subsets and the presence/absence of atherosclerotic plaques in ART-treated PLWH.

| Models |  | Fibrinogen | Memory<br>Th17/Treg<br>ratio | EM<br>Th17/Treg<br>ratio | CM<br>Th17/Treg<br>ratio | CCR9<sup>+</sup><br>HLA-DR<sup>low</sup><br>non-<br>classical<br>monocytes | CCR9<sup>low</sup><br>HLA-DR<sup>high</sup><br>non-<br>classical<br>monocytes | M-DC8+<br>HLA-DR<sup>high</sup><br>monocytes |
| --- | --- | --- | --- | --- | --- | --- | --- | --- |
| Crude Model | OR (CI) | 2.51<br>(1.22-5.18) | 0.49<br>(0.27-0.86) | 0.44<br>(0.24-0.79) | 0.57<br>(0.33-0.99) | 0.30<br>(0.15-0.63) | 3.56<br>(1.45-8.74) | 2.71<br>(1.24-5.91) |
|  | Adj. p | **0.01** | **0.03** | **0.03** | **0.05** | **0.01** | **0.02** | 0.10 |
| Model 1 | OR (CI) | 4.33<br>(1.35-14) | 0.47<br>(0.22-1.02) | 0.43<br>(0.18-1.03) | 0.55<br>(0.27-1.11) | 0.24<br>(0.09-0.62) | 6<br>(1.77-20.28) | 5.39<br>(1.56-18.6) |
|  | Adj. p | **0.03** | 0.1 | 0.1 | 0.1 | **0.015** | **0.015** | **0.04** |
|  | %Δ OR | 72% | 3% | 2% | 4% | 22% | 70% | 98% |
| Model 2 | OR (CI) | 3.87<br>(1.3-11.4) | 0.45<br>(0.21-0.94) | 0.41<br>(0.18-0.94) | 0.52<br>(0.26-1.03) | 0.2<br>(0.07-0.56) | 6.44<br>(1.89-21.95) | 6.06<br>(1.68-21.93) |
|  | Adj. p | **0.03** | **0.07** | **0.07** | **0.07** | **0.01** | **0.01** | **0.03** |
|  | %Δ OR | 54% | 9% | 6% | 9% | 35% | 80% | 123% |
| Model 3 | OR (CI) | 4<br>(1.4-12) | 0.41<br>(0.19-0.91) | 0.38<br>(0.16-0.94) | 0.48<br>(0.24-0.97) | 0.17<br>(0.06-0.52) | 8.09<br>(2.06-31.67) | 7.13<br>(1.8-28.22) |
|  | Adj. p | **0.02** | **0.06** | **0.06** | **0.06** | **0.01** | **0.01** | **0.03** |
|  | %Δ OR | 62% | 17% | 13% | 16% | 44% | 126% | 162% |
| Model 4 | OR (CI) | 3.37<br>(1.3-8.5) | 0.45<br>(0.23-0.89) | 0.47<br>(0.24-0.92) | 0.49<br>(0.26-0.94) | 0.26<br>(0.11-0.6) | 5<br>(1.72-14.56) | 3.75<br>(1.4-10.04) |
|  | Adj. p | **0.02** | **0.05** | **0.05** | **0.05** | **0.01** | **0.01** | **0.05** |
|  | %Δ OR | 34% | 7% | 6% | 14% | 15% | 40% | 40% |
| Fibrinogen | OR (CI) | N/A | 0.45<br>(0.24-0.86) | 0.45<br>(0.23-0.87) | 0.5<br>(0.27-0.94) | 0.26<br>(0.11-0.62) | 3.07<br>(1.14-8.25) | 2.75<br>(1.12-6.74) |
|  | Adj. p | N/A | **0.04** | **0.04** | **0.04** | **0.02** | 0.1 | 0.15 |
|  | %Δ OR | N/A | 8% | 3% | 13% | 13% | 14% | 1% |

OR, odds ratio; CI, 95% confidence interval; Adj. p, adjusted p-values; %Δ (delta) OR, OR difference relative to the crude model. FSR, Framingham score risk; N/A, not applicable
Model 1: ART duration, smoking and statins
Model 2: ART duration, smoking and LDL
Model 3: ART duration, smoking and triglycerides
Model 4: ART duration, HIV duration, FRS

## Results

### Laboratory and imaging markers of subclinical atherosclerosis in ART-treated PLWH

In previous studies performed by our group, CTA-Scan was used as a noninvasive tool to visualize and quantify CAA plaque as total (TPV) and low attenuated plaque volume (LAPV) in ART-treated PLWH (HIV^+^) and uninfected controls (HIV^−^) included in our CHACS cohort [5,55–58]. In the current cardiovascular imaging sub-study, a total of n=61 HIV^+^ and n=21 HIV^−^ were included, with demographic and clinical characteristics depicted in Table 1. Briefly, differences between HIV^+^ and HIV^−^ groups in terms of age (mean 55.38 *versus* 56.94 years), body mass index (BMI; median 24.43 *versus* 25.72 Kg/m^2^), and FRS (median 10 *versus* 8) did not reach statistical significance; also, similar proportions of participants received treatment with statins (23% *versus* 23.8%) (Table 1). The HIV^+^ group included 61/61 males (100%), while the HIV^−^ group included only 17/21 males (81%; p=0.003) (Table 1). The measurement of CAA plaque volume using CT-Scan angiograms was performed in all sub-study participants, as we previously reported [5,56]. The CAA plaque prevalence was similar between the HIV^+^ and HIV^−^ groups (Table 1). Indeed, among the HIV^+^ group, 39/61 (64%) participants were identified with detectable plaque (>1), while 22/61 (36%) participants had undetectable plaque (<1) measured as TPV and LAPV (plaque volume expressed in mm^3^) (Supplemental Figure 1A-B; Table 1). The HIV^−^ group included 13/21 (62%) with detectable plaque, while 8/21 (38%) participants had undetectable plaque (Supplemental Figure 1A-B; Table 1). Similarly, differences in the CAA plaque volume between HIV^+^ *versus* HIV^−^ groups did not reach statistical significance [TPV (median 108.3 *versus* 49.7) and LAPV values (median 34.4 *versus* 10.53); p=0.32 and 0.39, respectively] (Table 1).

In terms of laboratory parameters, the HIV^+^ and HIV^−^ groups presented with similar white blood counts, lymphocyte counts, and LDL levels; however, HIV^+^ compared to HIV^−^ groups had significantly lower levels of HDL (median: 1.23 *versus* 1.44) and higher levels of triglyceride (median: 1.73 *versus* 1.21) (Table 1). Since low HDL and high triglycerides are two well-documented CVD risk factors [61] such differences may accelerate the occurrence of subclinical CVD events in ART-treated PLWH, consistent with our recently reported findings [5].

We further sought to identify clinical and laboratory markers associated with subclinical CAA imaging among HIV^+^ participants (Table 2). Considering the fact that TPV and LAPV values were strongly positively correlated in HIV^+^ and HIV^−^ participants (Supplemental Figure 1C; p<0.0001, r=0.9913), all subsequent statistical analyses were based on the TPV values equal or higher than zero. In terms of demographics and clinical parameters, TPV^+^ and TPV^−^ HIV^+^ participants were similar in age (mean: 56.24 *versus* 53.85 years), sex (100% males) and the BMI (median: 24.05 *versus* 25.29 Kg/m^2^). The Framingham score (median 11 *versus* 8; p=0.084) and the number of HIV^+^ participants under statin treatment [(12/39; 30.8%) *versus* (2/22; 9.1%); p=0.064] tented to be superior among TPV^+^ compared to TPV^−^ participants (Table 2). Finally, the smoking was more prevalent in TPV^+^ *versus* TPV^−^ HIV^+^ participants (p=0.005) (Table 2).

In terms of HIV disease parameters, the frequency of HIV^+^ participants with undetectable plasma viral loads was not statistically different in TPV^+^ compared to TPV^−^ groups (94.8 *versus* 86.36%, p=0.34). However, TPV^+^ compared to TPV^−^ HIV^+^ participants were on ART for a longer time (mean 16.14 *versus* 10.69; p=0.003) and the time since infection tended to be superior (median: 19.6 *versus* 14.7; p=0.06) (Table 2). Nadir CD4 counts were similar in TPV^+^ compared to TPV^−^ groups (median 195 *versus* 170 cells/µl; p=0.397). Regarding laboratory parameters, TPV^+^ and TPV^−^ HIV^+^ participants showed similar white blood cells and lymphocyte counts, and D-dimer, LDL, HDL, and triglycerides levels; however, fibrinogen levels, indicative of coagulopathy [62], were significantly increased in the TPV^+^ compared to TPV^−^ HIV^+^ (Table 2; p=0.005). Thus, subclinical signs of CVD reflected by the presence of CAA plaque were strongly associated with a longer time on ART, as well as increased plasma fibrinogen levels.

### Plasma markers in relationship with HIV-1 status and subclinical atherosclerosis

To identify systemic biomarkers associated with subclinical atherosclerosis, we quantified plasma levels of well-established markers of gut damage (sCD14, LBP, FABP2) and systemic inflammation (CCL20/MIP-3α, MIF, CX3CL1/FKN, CCL25/TECK) in the HIV^−^ and HIV^+^ groups, and in the HIV^+^ group in relationship with subclinical CVD. While higher plasma levels of sCD14, FABP2, CCL20, MIF and CX3CL1 distinguished HIV^+^ from HIV^−^ participants (Supplemental Figure 2A-B), only levels of CCL20, previously associated with HIV disease progression [63], were significantly increased in TPV^+^ compared to TPV^−^ among HIV^+^ participants (p=0.0301) (Supplemental Figure 2C-D). Thus, plasma CCL20 levels were increased in ART-treated PLWH with subclinical CVD.

### T-cell profile alterations in relationship with HIV-1 status and subclinical atherosclerosis

To study the relationship between key players in adaptive immunity and subclinical CVD status, polychromatic flow cytometry was used to identify specific T-cells subsets in the peripheral blood of HIV^+^ and HIV^−^ participants with/without CAA (Supplemental Figure 3). Staining with CD45RA and CCR7 was used to identify memory subsets, as we previously reported [64]. Surface staining with CD25 and CD127 Abs and intra-nuclear staining with FOXP3 Abs allowed the identification of regulatory T cells (Tregs; CD127^low^CD25^high^FoxP3^+^ phenotype), as we previously reported [64]. Finally, Th17-polarized T-cells were identified among FOXP3^−^ cells by Boolean combination of gates between CCR6, CD26 and CD161 and defined as cells with a CCR6^+^CD26^+^CD161^+^ phenotype, as previously reported by our group and others [64,65].

HIV^+^ compared to HIV^−^ participants showed decreased frequencies of CD4^+^ T-cells within CD3^+^ T-cells, with increased frequencies of CD8^+^ T-cells and lower CD4/CD8 ratios (Supplemental Figure 4A), consistent with previous literature identifying the CD4/CD8 ratio as a marker of HIV disease progression [66] and viral reservoirs [67,68]. No statistically significant differences in the latter parameters were observed among TPV^+^ and TPV^−^ HIV^+^ participants (Supplemental Figure 4B). The frequencies of total memory (CD45RA^−^) CD4^+^ T-cells, as well as those of effector memory (EM, CD45RA^−^CCR7^−^), central memory (CD45RA^−^CCR7^+^; CM), and EM expressing CD45RA (EMRA) CD4^+^ T-cells, did not differ between HIV^+^ and HIV^−^ participants (Supplemental Figure 5A), neither between TPV^+^ and TPV^−^ HIV^+^ participants (Supplemental Figure 5B). Of notice, HIV^+^ compared to HIV^−^ participants showed a statistically significant increase in the frequency of Tregs (p=0.0002), similar frequencies of CD4^+^ T-cells with a Th17 phenotype within total memory, EM and CM subsets, and reduced total memory Th17/Treg (p=0.029), EM Th17/Tregs (p=0.007) and CM Th17/Tregs (p=0.047) ratios (Supplemental Figure 6A-C).

Among HIV^+^ participants, TPV^+^ compared to TPV^−^ individuals exhibited similar frequencies of Tregs, but decreased frequencies of total memory (p=0.0166), EM (p=0.0083) and CM (p=0.0149) subsets with a Th17 phenotype, and decreased Th17/Treg ratios among memory (p=0.0373) and EM (p=0.0259) subsets (Figure 1A-C). The same trend was not observed in HIV^−^ individuals, although the low number of HIV^−^ participants available for the study may limit valid interpretations (Supplemental Figure 7A-C). Thus, ART-treated HIV-1 infection is associated with an expansion of Tregs and an alteration of Th17/Treg ratios, with the paucity of Th17 cells and the subsequent impact on Th17/Treg ratios being exacerbated in HIV^+^ participants with subclinical CAA.

**Figure 1:**
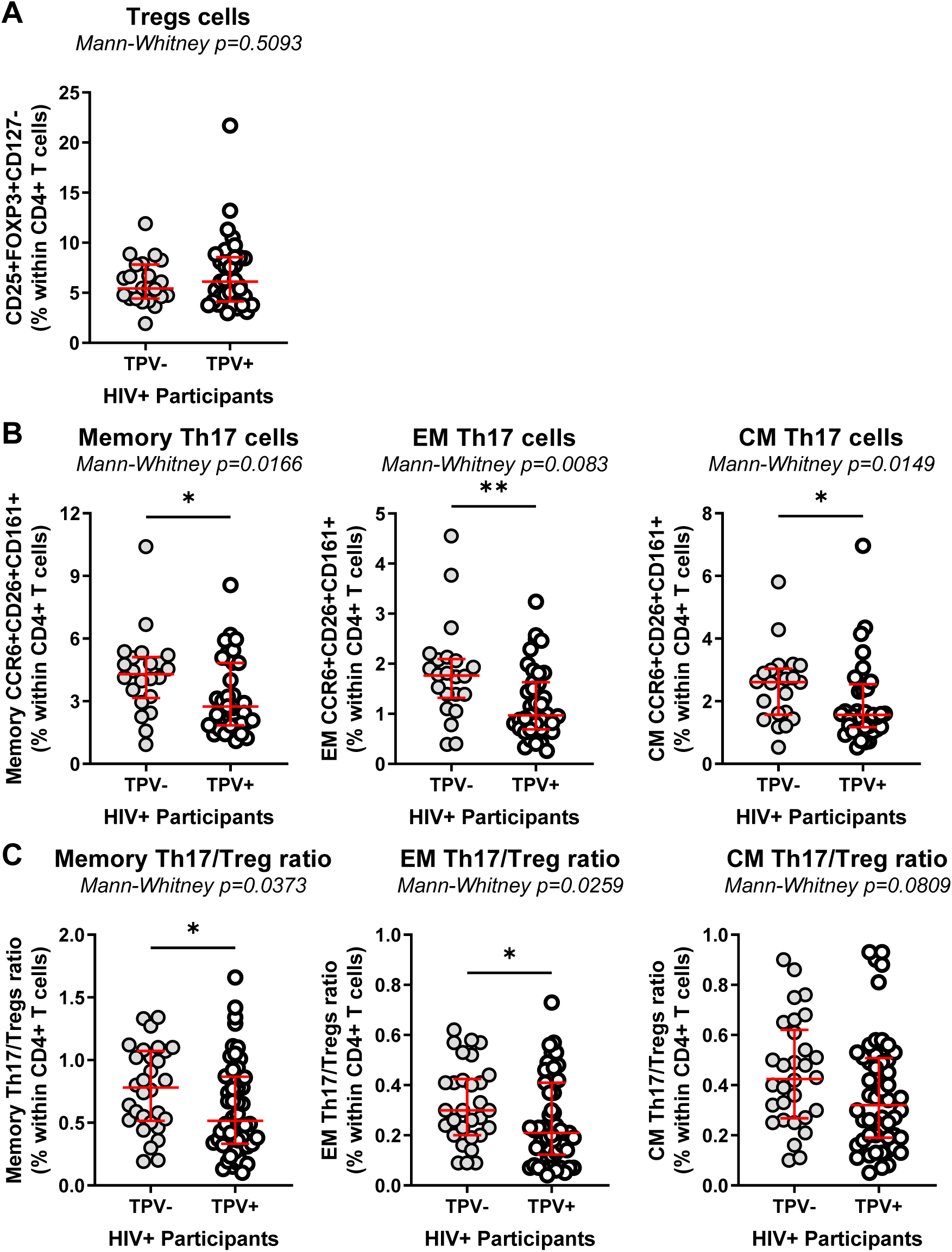
Decreased Th17 frequencies and Th17/Tregs ratios coincide with subclinical CAA in ART-treated PLWH. The frequencies of regulatory CD4^+^ T-cells (Tregs; CD25^+^FOXP3^+^CD127^−^CD4^+^) **(A)**, as well as Th17-polarized CD4^+^ T-cells (CCR6^+^CD26^+^CD161^+^) with memory (CD45RA^−^), effector memory (EM) (CD45RA^−^CCR7^−^) and central memory (CM) (CD45RA^−^CCR7^+^) phenotypes **(B)**, were compared between TPV^+^ (n=39) and TPV^−^ (n=22) HIV^+^ participants. **(C)** Shown are the Th17/Treg ratios within the memory, EM and CM Th17 subsets among TPV^+^ *versus* TPV^−^ HIV^+^ participants. Median and IQ range are indicated in red as horizontal lines. Differences among study groups were determined by Mann-Whitney rank test. P-values and statistical significance are indicated in the figures (*, p < 0.05; **, P < 0.01; ***, P < 0.001).

### Monocyte alterations in relationship with HIV-1 status and subclinical atherosclerosis

Changes in monocyte heterogeneity have been associated with various pathological conditions, including HIV-1 infection [69]. Thus, we investigated changes in monocyte subset frequencies and their phenotypes in relationship with the HIV and TPV status. The flow cytometry gating strategy used to identify monocyte subsets is depicted in Supplemental Figure 8, as previously described [34,64,70,71]. An expansion of classical (CD14^++^CD16^−^) (p=0.036) to the detriment of intermediate (CD14^++^CD16^+^) (p=0.039) monocytes was observed in HIV^+^ compared to HIV^−^, whereas the frequencies of non-classical (CD14^−^CD16^++^) and M-DC8^+^ (Slan^+^) monocytes were similar (Supplemental Figure 9A). Within the HIV^+^ group, classical, intermediate, non-classical and Slan monocytes were similar in frequency regardless of the TPV status (Supplemental Figure 9B).

The phenotype of monocyte subset was further analyzed, including the expression of the chemokine receptors CCR2 (involved in monocyte trafficking from bone marrow into peripheral tissues [72], CX3CR1 (involved in intermediate and non-classical monocyte patrolling onto vascular beds[73], and CCR9 (involved in cell recruitment into multiple sites including the atherosclerotic plaque [74]. Consistent with previous reports [75–77], CCR2 and CCR9 were mainly expressed on classical monocytes, while CX3CR1 was predominantly expressed on non-classical and M-DC8^+^ monocytes (Figure 2; Supplemental Figure 10). The expression of CCR2 and CX3CR1 on monocyte subsets did not differ between HIV^+^ and HIV^−^ groups (Supplemental Figure 10A-D). In contrast, the expression of CCR9 on intermediate and non-classical monocytes was significantly diminished in HIV^+^ compared to HIV^−^ groups (p=0.0087 and p=0.0066, respectively) (Supplemental Figure 10B-C). A statistically significant increase in the expression of HLA-DR was observed on non-classical (p=0.0258) and Slan/M-DC8^+^ monocytes (p=0.0384) of HIV^+^ compared to HIV^−^ (Supplemental Figure 10C-D). Further, the stratification based on TPV in HIV^+^ participants demonstrated similar expression of CCR2 and CX3CR1 on all monocyte subsets, a decreased expression of CCR9 on non-classical monocytes (p=0.0024), and an increased expression of HLA-DR on non-classical (p=0.0081) and Slan/M-DC8^+^ (p=0.0091) monocytes from TPV^+^ compared to TPV^−^ HIV^+^ participants (Figure 2A-D). Thus, in the cohort used for this study, significant alterations were observed in terms of frequency of classical and intermediate monocyte subsets in relationship with the HIV but not the TPV status; however, changes were observed in the expression of HLA-DR and CCR9, with the predominance of CCR9^low^HLA-DR^high^ non-classical and HLA-DR^high^ M-DC8^+^ monocytes in HIV^+^ *versus* HIV-participants, a difference further exacerbated within the HIV^+^ group in TPV^+^ *versus* TPV^−^ participants.

**Figure 2.**
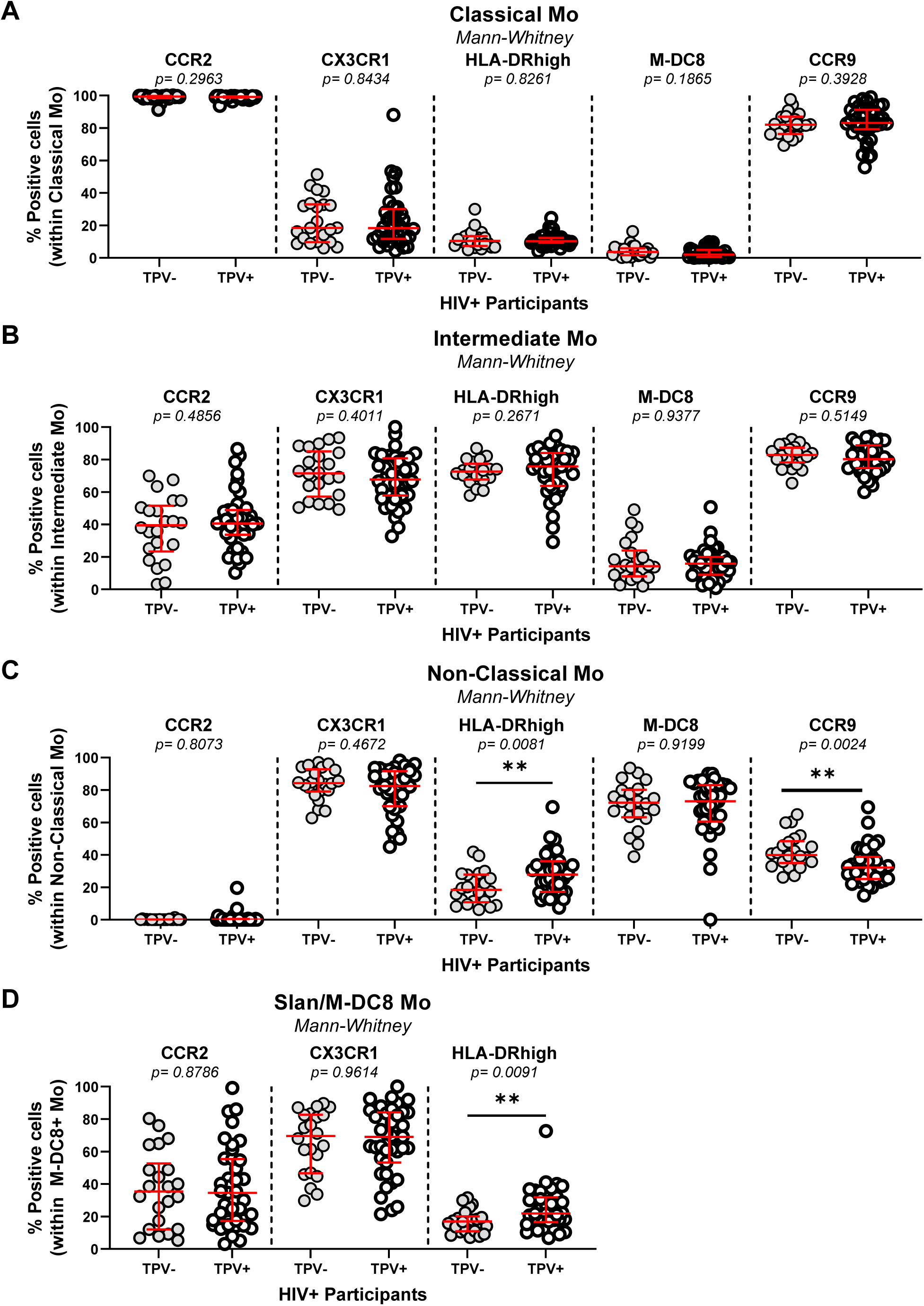
Non-classical CCR9^low^HLA-DR^high^ and M-DC8^+^HLA-DR^high^ monocytes are expanded in ART-treated PLWH with subclinical CAA. Total monocytes (Mo) were identified as cells lacking the T cell lineage markers CD3 and CD4 and the DC marker CD1c, and expressing HLA-DR (Supplemental Figure 7). Shown is the expression of CCR2, CX3CR1, CCR9, HLA-DR and M-DC8 on classical (CD14^++^CD16^−^) **(A)** intermediate (CD14^++^CD16^+^) **(B)**, and non-classical (CD14^+^CD16^++^) Mo **(C)**, as well as the expression of CCR2, CX3CR1 and HLA-DR on M-DC8+ Mo **(D),** in PBMC of TPV^+^ (n=39) *versus* TPV^−^ (n=22) HIV^+^ participants. Median and IQ range are indicated in red as horizontal lines. Differences among study groups were determined by Mann-Whitney rank test. P-values and statistical significance are indicated in the figures (*, p < 0.05; **, P < 0.01; ***, P < 0.001).

### mDC and pDC alterations in relationship with HIV-1 status and subclinical atherosclerosis

DC are important immune players acting at the interface between innate and adaptive immunity. While myeloid DC (mDC) are mainly involved in antigen-presentation and priming the T cell response [78,79], plasmacytoid DC (pDC) are specialized in sensing viruses and producing type I IFN [78]. The flow cytometry gating strategy used to identify mDC (HLA-DR^+^CD1c^+^) and pDC (BDCA2^+^CD123^+^) is depicted in Supplemental Figure 11. Briefly, mDC and pDC were gated on CD3^−^CD4^−^ Live cells and the expression of CCR6 and CCR9 was also evaluated in each subset. The frequency of mDC and their expression of CCR6 or CCR9 was similar in HIV^+^ and HIV^−^ participants (Supplemental Figure 12A), with no differences between TPV^+^ and TPV^−^ HIV^+^ participants (Supplemental Figure 12B). Similar results were obtained for the frequency and phenotype of pDC (Supplemental Figure 13A-B). Thus, in this cohort, the mDC and pDC frequencies and CCR6/CCR9 phenotype did not distinguish between groups with different HIV and TPV status.

### Multivariate analysis identifies an immunological signature associated with subclinical atherosclerosis in ART-treated PLWH

A logistic regression model was used to determine the association between covariates and the presence or the absence of subclinical CAA plaque (*i.e.,* TPV). In crude analysis, fibrinogen levels, the Th17/Treg ratios among total memory, EM, and CM CD4^+^ T-cell subsets, and the frequency of CCR9^low^HLA-DR^high^ non-classical monocytes were associate with the presence of CAA plaque, while the frequency of CCR9^+^HLA-DR^low^ non-classical monocytes was associated with the absence of CAA plaque (Table 3). Adjustments were performed for ART duration, smoking and stains (Model 1), ART duration, smoking and LDL (Model 2), ART duration, smoking and triglycerides (Model 3), and ART duration, HIV duration and FRS (Model 4), as well as for FRS and fibrinogen separately. The association remained statistically significant for fibrinogen, and CCR9^low^HLA-DR^high^ and CCR9^+^HLA-DR^low^ non-classical monocytes, in all Models 1-4 and for Th17/Treg ratios in Model 4 only (Table 3). Interestingly, M-DC8^+^ monocytes proved to be associated with the presence of TPV only after the adjustment in Models 1-4 (Table 3). Finally, after adjustment with fibrinogen alone, total memory, EM and CM Th17/Treg ratios, remained associated with the presence of plaque, while the frequency of CCR9^+^HLA-DR^low^ non-classical monocytes remained associated with the absence of plaque (Table 3).

Together, our studies reveal alterations in Th17/Treg ratios and non-classical monocyte frequency/phenotype that, together with fibrinogen levels, coincide with subclinical CVD, alterations that may directly/indirectly fuel CAA plaque formation in ART-treated PLWH.

## Discussion

Previous studies by our group and others documented immunological alterations that occur in PLWH and persist during ART, including gut barrier impairment, microbial translocation, Th17 cell paucity, and monocyte subset expansion/activation (reviewed in [69,80–82]). In this manuscript, we investigated whether these alterations are associated with subclinical CAA in ART-treated PLWH in our local Canadian cohort mainly composed of male participants. To this goal, we had access to plasma and PBMC samples from ART-treated PLWH (HIV^+^) and HIV^−^ participants included in the cardiovascular imaging sub-study of the CHACS [54–56]. Markers of gut dysfunction (*i.e.,* sCD14, LBP, I-FABP) and chemokines involved in cell trafficking (*i.e.,* CCL20, CX3CL1, CCL25) were quantified in the plasma, while the frequency and phenotype of CD4^+^ T-cell subsets (*i.e.,* Th17, Tregs), CD8^+^ T-cells, as well as monocyte (*i.e.,* classical, intermediate, non-classical, M-DC8^+^) and DC (*i.e.,* mDC, pDC) subsets were monitored in the peripheral blood. Variations in these immunological parameters, together with multiple clinical measurements, were studied in relationship with the presence of CAA plaque visualized/monitored by CT Scan, as we previously reported [5,56–58]. Our results support a model in which the paucity of Th17 cells, the alteration of Th17/Treg ratios, and the abundance of CCR9^low^HLA-DR^high^ non-classical monocytes favor a state of systemic immune activation that may fuel CAA in ART-treated PLWH.

Studies previously performed by our group on a total of 181 ART-treated PLWH (HIV^+^) and 84 HIV-uninfected controls (HIV^−^) included in the CHACS at baseline, revealed a two to three-fold increase in the coronary non-calcified plaque burden in ART-treated PLWH compared to HIV^−^ individuals [5]. The current immunological sub-study was performed on a fraction of HIV^+^ (n=61) and HIV^−^ (n=21) participants, the first participants enrolled in the CHACS [54–56]. In this sub-study, statistical differences were not observed in the prevalence of CAA plaque detection (TPV >0) between HIV^+^ and HIV^−^ groups. Nevertheless, the number of participants with TPV values >100 mm^3^ was higher in HIV^+^ (32/61; 52.4%; median TPV: 608.7 mm^3^) compared to HIV^−^ (7/21; 33.3%; median TPV: 192.8 mm^3^). Consistently, we observed that levels of HDL were lower, while the levels of triglycerides were higher in ART-treated PLWH compared to HIV^−^ participants. In addition, plasma markers of intestinal damage (FABP2), systemic inflammation (sCD14), and chemokines involved in cell trafficking (CCL20, MIF, and CX3CL1) were significantly higher in HIV^+^ compared to HIV^−^ participants. This is indicative that ART does not restore gut barrier functions, nor counteracts systemic immune activation in PLWH, thus explaining an increased CVD risk associated with HIV-1 infection relative to the general population [7,8,10,13–15].

The detection of subclinical CAA plaque in HIV^+^, as measured by TPV and LAPV, was associated with a longer time on ART and elevated plasma levels of fibrinogen, as well as a tendency for an increased time since infection, Framingham score, and the use of statins. No differences in nadir CD4 counts were observed. Among soluble factors monitored in plasma, only levels of the CCR6 binding chemokine CCL20 proved to be higher in TPV^+^ compared to TPV^−^ HIV^+^. CCL20 was previously reported to be increased during HIV infection [63]. Studies in CCR6^-/-^Apo^-/-^ mice demonstrated the role of CCR6 in monocyte recruitment into atherosclerotic plaques, as well as monocyte emigration from bone marrow [55,83]. Of note, monocyte *per se* can produce CCL20 upon exposure to HIV [84]. Thus, increased plasma levels of CCL20 point to an increases risk of CVD in ART-treated PLWH.

Th17-polarized CD4^+^ T-cells represent keystone players of immunity at mucosal barriers [80–82]. Multiple studies documented alterations in Th17 function and frequency during HIV/SIV infection and identified Th17 depletion as the major cause for gut barrier damage, which results in microbial translocation-induced systemic immune activation [80–82]. An important finding of our study is represented by the significant reduction in the frequency of Th17 cells with memory, EM and CM phenotypes in the peripheral blood of TPV^+^ compared to TPV^−^ HIV^+^ participants. The preferential depletion of Th17 cells during HIV-1 infection was linked to their relatively high susceptibility to integrative HIV-1 infection [80–82]. The loss of Th17 cells during early acute HIV infection is correlated with systemic immune activation measured by the proportion of activated CD8^+^ T-cells [85]. Additionally, higher plasma lipopolysaccharide, which is a marker of microbial translocation and gut damage, correlated with lower frequencies of Th17 cells in chronic HIV infection [86]. ART initiation during very early acute HIV infection restored Th17 cells. It was also reported that although long-term ART adherence normalizes the frequency of sigmoid Th17 cells, high levels of plasma lipopolysaccharide were still observed in those individuals [86,87]. In our cohort, we did not observe differences in the frequencies of Th17 cells between HIV^−^ and HIV^+^ participants. This similarity could be explained by the fact that the median ART duration of our participants was relatively high, exactly 15.67 years (IQR: 8.14-19.37 years), raising the possibility that long-term ART may have restored the Th17 compartment in the blood. Nevertheless, the detection of CAA plaque in ART-treated PLWH was associated with a reduced frequency of Th17 cells, which may mirror alterations in mucosal barrier integrity. The existence of a link between alterations occurring at the level of the mucosal intestinal barrier and the pathology of the cardiovascular system supports the existence of the gut-heart axis [10,17,22,31,33,43,46,55,61,80,87–93].

The detection of IL-17A-producing CD4^+^ T-cells in the atherosclerotic plaque of mice [2–4,6,13,18,20,28,33,81,94–96] supports the possibility that subsets of Th17 cells may be recruited into the vascular beds and locally fuel atherosclerosis. In line with this scenario, in our study TPV^+^ compared to TPV^−^ ART-treated PLWH exhibit increased plasma levels of CCL20, a chemokine essential for CCR6^+^ Th17 cell trafficking [80–82]. Considering that T-cell functions depend on TCR triggering, future studies should determine the antigenic specificity of Th17 cells recruited into the atherosclerotic plaques. Indeed, studies reported that CD4^+^ T-cells infiltrating the atherosclerotic plaque recognize CMV and HIV peptides, as well as LDL and apolipopotein B peptides [24]. Th17 cell mainly recognize components of the microbiota [97,98]. Or, specific components of the microbiota were reported to be present into atherosclerotic plaques [99,100], thus supporting a cognate contribution of Th17 cells to the process of atherosclerosis. Nevertheless, the role of Th17 cells may be dual, a positive one at the intestinal level, where they maintain the integrity of the mucosal barrier, and a deleterious one when infected by HIV and recruited into the atherosclerotic plaque and exerting effector functions in response to translocated microbial components.

In contrast to Th17 cells, Tregs are reported to be expanded during HIV-1 infection despite the initiation of ART, with their suppressive functions being detrimental for the proper development of anti-HIV immunity [101]. Due to alterations in the frequency of Th17 and Tregs during HIV-1 infection, the Th17/Tregs ratio is decreased during HIV infection [102,103]. In primary HIV-1 infection, the Th17/Tregs ratio was negatively correlated with the proportion of activated CD8^+^ T-cells, plasma viral load, and markers of monocyte activation such as sCD14 and IL-1RA [103]. In line with these results, we observed increased frequencies of Tregs and decreased Th17/Tregs ratios in this group of ART-treated PLWH compared to HIV^−^ participants included in the CHACS. Although we did not observe statistically significant differences in the frequency of Tregs between TPV^+^ and TPV^−^ in our group of n=61 ART-treated PLWH, another sub-study performed by our group on n=84 participants from the same CHACS cohort found an increase in Treg frequency in TPV^+^ compared to HIV^+^ participants [27]. Despite this discrepancy that may be explained by differences in the number of participants included in the two studies, we consistently report here that CAA is associated with a reduction in Th17/Treg ratios in ART-treated PLWH.

Solid literature evidence supports the contribution of specific monocyte subsets to the process of atherosclerosis [34,35], with studies in mice demonstrating the recruitment of non-classical monocytes into the plaque *via* the chemokine receptor CX3CR1 and its ligand CX3CL1/FKN [104]. Non-classical monocytes indeed express CX3CR1 preferentially and patrol the vascular beds rich in membrane-associated CX3CL1 (reviewed in [69]). Of note, an increased proportion of non-classical monocytes expressing CX3CR1 were positively associated with coronary intima-media thickness [42,105]. Non-classical monocytes also produce inflammatory cytokines (IL-6), chemokines (CCL2), and MMP-9 upon interaction with CX3CL1-expressing endothelial cells [106]. In addition, the CCL2 receptor CCR2 is expressed on classical and intermediate monocytes and is key in establishing atherosclerosis in mice models [34,35]. In our study, although we did not observe an expansion of intermediate/non-classical CCR2^−^CX3CR1^+^ monocytes, subclinical CAA was associated with the abundance of CCR9^low^HLA-DR^high^ non-classical monocytes. The significance of this correlation was maintained after cofounder factor adjustment and multivariate analyses, thus raising new questions on the role of these monocytes in CVD in ART-treated PLWH.

The expression of CCR9 on monocytes and the CCR9/CCL25 axis have been associated with autoimmune diseases such as inflammatory bowel disease, rheumatoid arthritis (RA) and ankylosing spondylitis [76,107]. In relation to CVD, in a mouse model of atherosclerosis, the presence of monocytes and resident macrophages expressing CCR9 with characteristics of plaque-foaming cells was observed. In addition, CCR9 silencing by RNA interference decreased the size of atherosclerotic plaques [74]. CCR9 is also associated with cardiac hypertrophy in mice models [108]. Furthermore, in humans, CCR9^+^ cells were visualized infiltrating the atherosclerotic plaques [74]. In our cohort, we observed a decreased expression of CCR9 on non-classical monocytes in ART-treated PLWH with subclinical CAA. One of the possible explanations is the infiltration of CCR9^+^ non-classical monocytes into the atherosclerotic plaques, explaining why they are not observed in the periphery. On the contrary, CCR9^+^ pDCs have been reported to be tolerogenic and CCR9 loss to be deleterious in the development of atherosclerosis [51], which suggests that CCR9 is not necessarily a marker of inflammation. In our study, the predominance of CCR9^+^ non-classical monocytes was associated with the absence of CAA plaque. Therefore, one cannot exclude the possibility that CCR9^+^ monocytes play a protective role in atherosclerosis. Changes in CCR9 expression did not correlate with changes in plasma levels of the CCR9 ligand CCL25. Future mechanistic studies should address this possibility.

Our study has multiple limitations. First, we performed cross-sectional but not longitudinal immunological studies on baseline samples included in the CHACS cohort. This pan-Canadian cohort was initiated in Montréal in 2013 and includes now n=805 HIV^+^ and n=244 HIV^−^ controls for a 5-year longitudinal follow-up [54–56]. Participants in the CHACS cardiovascular imaging sub-study (n=181 HIV^+^ and n=84 HIV^−^) are presently undergoing repeat Cardiac CT angiogram, which will grant data to inform us on the validity of the identified markers of subclinical CAA in their ability to predict CVD progression in ART-treated PLWH. The current results will orient future longitudinal studies on the CHACS cohort. Second, the CHACS participants are in majority males, and sex-related differences could not be addressed. Indeed, it is reported that differences exist between males and females in terms of HIV and CVD pathogenesis [109], with an increased CVD risk observed in female *versus* male PLWH [3,95]. Whether the immunological signature we identified is also associated with subclinical CVD in women remains to be investigated. Third, we did not study alterations in the Th17/Tregs ratios in relationship with the size of HIV reservoirs. However, studies performed by our group on the CHACS samples revealed an increased HIV-DNA reservoir in CD4^+^ T-cells of TPV^+^ *versus* TPV^−^ HIV^+^ [96], pointing to the possibility that Th17 depletion is a consequence of their infection. Fourth, we were not able to confidently conclude on immunological differences between HIV^−^ participants with and without subclinical CAA due to the reduced number of individuals enrolled at the time we performed the study. Nevertheless, despite the small sample size, there was no decrease in Th17 frequency nor in Th17/Treg ratios between TPV^+^ and TPV-HIV^−^ participants, indicative that the immunological signature associated with subclinical CAA is likely distinct in HIV^+^ *versus* HIV^−^ groups.

In conclusion, we report results generated on peripheral blood cells collected from participants included in the CHACS, a longitudinal cohort established to identify novel biomarkers predictors of subclinical CVD in ART-treated PLWH [54–56]. Our investigations revealed an immunological signature associated with subclinical CAA in ART-treated PLWH, including ***i)*** increased plasma CCL20 levels (indicative of systemic immune activation); ***ii)*** reduced Th17/Treg ratios (reflecting ongoing alterations of mucosal barrier integrity); ***iii)*** increased frequencies of non-classical CD16^+^ monocytes with a CCR9^low^HLA-DR^high^ phenotype (exerting pro-inflammatory features), and ***iv)*** increased plasma fibrinogen levels (sign of coagulopathy). Our results reveal immune cell subsets that may predict CAA in ART-treated PLWH and emphasize the need for new therapeutic strategies designed to restore immunological competence in PLWH under long-term ART, especially at mucosal barrier levels, in an effort to reduce the CVD risk.

## Authors’ contributors

TRWS contributed to study design, performed most of the experiments, analyzed results, prepared figures, and wrote the manuscript. YZ contributed to study design and performed experiments. AG and NFDR contributed to experiments. MEF was instrumental to the study design and provided protocols. AF performed all multivariate statistical analysis. JPR contributed to participant recruitment. CCL performed CT-SCAN angiographies and interpreted the results in all participants. ALL, MD and CLT contributed to patient recruitment, establishment of the biobank of plasma and PBMC, and provided access and valuable guidance in the analysis of clinical information, as well as statistical analysis. PA designed the study, analyzed results, contributed to figure preparation, and wrote the manuscript. All authors reviewed and approved the manuscript.

## Acknowledgments

This work was supported, in part, by funds from the Canadian Institutes of Health Research to PA (#MOP-114957; #TCO125276; IBC-154053), National Institutes of Health (NIH) to CT and PA (R01AG054324), as well as infrastructure funding from the Canadian Foundation for Innovation (CFI) to PA and CT. The *Fondation du CHUM* and the *Fonds de recherche du Québec – Santé* (FRQ-S) HIV/AIDS and Infectious Diseases Network supported core facilities and human cohorts. TRWS was supported by Doctoral awards from the Université de Montréal and the FRQ-S. The funding agencies were not involved in patient recruitment, data collection nor the analysis and interpretation of the results; and had no role in the writing of the manuscript nor the decision to submit it for publication.

The authors thank Dr. Dominique Gauchat and Philippe St Onge (Flow Cytometry Core Facility, CHUM-Research Center, Montréal, QC, Canada) for expert technical support with polychromatic flow cytometry sorting; Olfa Debbeche (NLC3 Core Facility CHUM-Research Center, Montréal, QC, Canada); and Mohamed Sylla and Etiene Larouche-Anctil for blood processing and management of cell biobank. We would also like to thank Mario Legault, Dr. Annie Chamberland, Stéphanie Matte, and Daniel Tremblay-Sher for their help with administrative duties and cohort database. The authors address a special thanks to all study participants for their crucial contribution to this work. J.-P.R is the holder of the Louis Lowenstein Chair in Hematology and Oncology, McGill University. CT holds the Pfizer/Université de Montréal Chair in HIV translational Research.

## Data sharing

Data from this manuscript is available from the corresponding author upon reasonable request.

