## Supplementary material for "A Blood Immunological Signature of Subclinical Coronary Artery Atherosclerosis in People Living with HIV-1 Receiving Antiretroviral Therapy": Suppl Table 1

### Supplemental Table 1: Antibodies and dyes used for polychromatic flow cytometry

| Target | Clone | Fluorochrome | Supplier |
|---|---|---|---|
| **Th17/Treg cocktail** | | | |
| CD3 | UCHT1 | Alexa Fluor 700 | Biolegend |
| CD4 | RPA-T4 | PerCP/Cyanine5.5 | Biolegend |
| CD45RA | HI100 | APC-eFluor 780 | eBioscience |
| CCR6 | 11A9 | PE | BD Pharmingen |
| CCR7 | 3D12 | BV786 | BD Horizon |
| CD25 | M-A251 | BV421 | BD Horizon |
| CD127 | HIL-7R-M21 | PE-Cy7 | BD Pharmingen |
| CD26 | M-A261 | APC | BD Pharmingen™ |
| CD161 | DX12 | BV650 | BD Horizon |
| FOXP3 | PCH101 | Alexa Fluor 488 | eBioscience |
| LIVE/DEAD™ Fixable Aqua Dead Cell Stain | | | Thermo Fisher Scientific |
| **Monocyte cocktail** | | | |
| CD3 | UCHT1 | Alexa Fluor 700 | Biolegend |
| CD4 | RPA-T4 | PerCP/Cyanine5.5 | Biolegend |
| CD1c | F10/21A3 | BV786 | BD OptiBuild |
| CD14 | M5E2 | APC | BD Pharmingen |
| CD16 | 3G8 | PE-Cy7 | BD Pharmingen |
| HLA-DR | L243 | APC/Cyanine7 | Biolegend |
| CCR6 | 11A9 | PE | BD Pharmingen |
| M-DC8/Slan | DD-1 | FITC | Miltenyi |
| CCR2 | K036C2 | PE/Dazzle 594 | Biolegend |
| CX3CR1 | 2A9-1 | BV421 | BD Horizon |
| LIVE/DEAD™ Fixable Aqua Dead Cell Stain | | | Thermo Fisher Scientific |
| **Monocyte and DC cocktail** | | | |
| CD3 | UCHT1 | Alexa Fluor 700 | Biolegend |
| CD4 | RPA-T4 | PerCP/Cyanine5.5 | Biolegend |
| CD1c | F10/21A3 | BV786 | BD OptiBuild |
| CD14 | M5E2 | APC | BD Pharmingen |
| CD16 | 3G8 | PE-Cy7 | BD Pharmingen |
| HLA-DR | L243 | APC/Cyanine7 | Biolegend |
| CCR6 | 11A9 | BV421 | BD Horizon |
| CCR9 | 112509 | PE | BD Pharmingen |
| CD123 | 6H6 | PE/Dazzle 594 | eBioscience |
| CD303 (BDCA2) | AC144 | FITC | Miltenyi |
| LIVE/DEAD™ Fixable Aqua Dead Cell Stain | | | Thermo Fisher Scientific |
