## Supplementary material for "A Blood Immunological Signature of Subclinical Coronary Artery Atherosclerosis in People Living with HIV-1 Receiving Antiretroviral Therapy": Suppl Figures 1-13

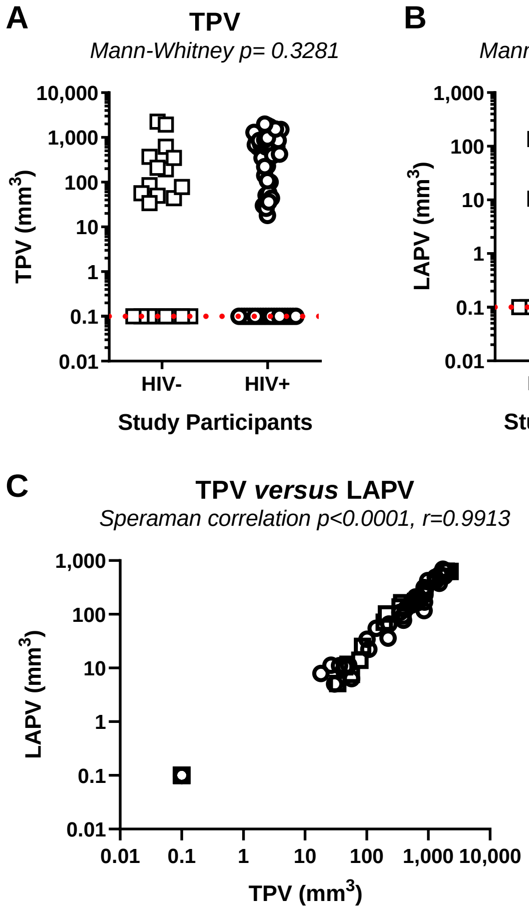


**Figure S1: Subclinical atherosclerosis in HIV-uninfected and ART-treated PLH participants.** A slice computed tomography (CT) scanner was used to perform noncontrast cardiac CT and coronary computed tomography angiography to measure **(A)** total plaque volume (TPV) and **(B)** low attenuated plaque volume (LAPV) in a cohort of asymptomatic CVD risk-matched HIV-uninfected (HIV-; n=21) and ART-treated PLH (HIV+; n=61) participants. Differences among groups were determined by Mann-Whitney rank test. P-values and statistical significance are indicated in the Figures. **(C)** The correlation between TPV and LAPV in $HIV^+$ (open symbols) and $HIV^-$ (grey symbols) participants was studied using the Spearman correlation model, with p and r values indicated on the graphs.

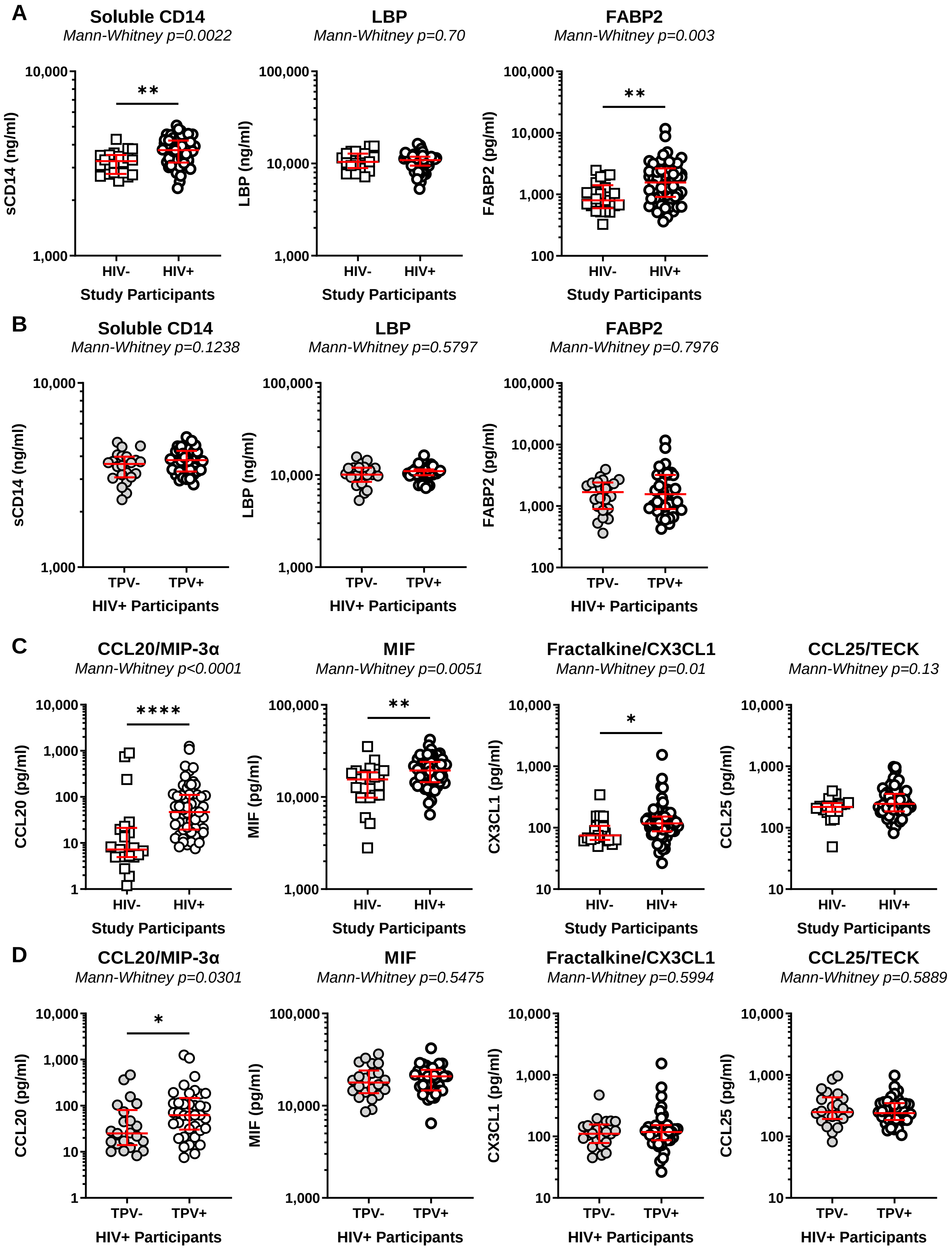


Wiche Salinas et al., Suppl. Figure 2

**Figure S2: Serological markers of microbial translocation, mucosal damage, and chemokines in HIV$^-$ and HIV$^+$ individuals with or without coronary plaque.** Markers of microbial translocation and gut damage (sCD14, LBP and FABP2; **A-B**) as well as markers of systemic inflammation (CCL20/MIP-3$\alpha$, MIF, CX3CL1/FKN, CCL25/TECK; **C-D**) were quantified in the plasma of HIV$^+$ (n=61) *versus* HIV$^-$ (n=21) participants (**A and C**) and also in the plasma of HIV$^+$ participants with (TPV$^+$; n=39) and without subclinical signs of atherosclerosis (TPV$^-$; n=22) (**B and D**). Median and IQ range are indicated in red as horizontal lines. Differences among study groups were determined by Mann-Whitney rank test. P-values and statistical significance are indicated in the figures (*, $p < 0.05$; **, $P < 0.01$; ***, $P < 0.001$).

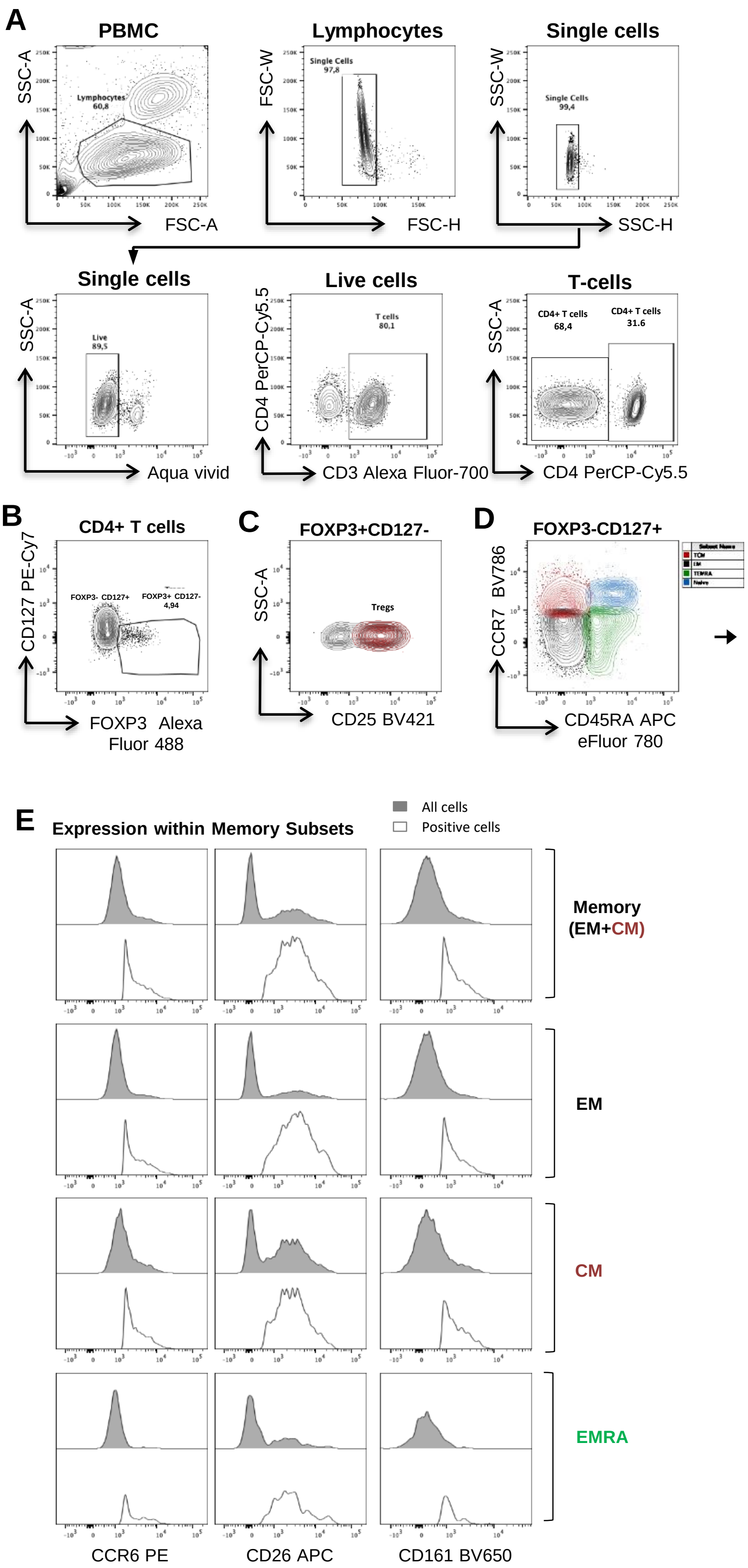


**Figure S3: Gating strategy for the identification of CD4⁺ T-cell subsets.** PBMCs from HIV⁺ and HIV⁻ participants were stained with fluorochrome-conjugated CD3, CD4, CD45RA, CCR7 CD25, CD127, CCR6, CD26, CD161, and FOXP3 Abs. Shown is the gating strategy to identify different T-cell subsets. Briefly, doublets and dead cells were excluded based on the FSC-W/FSC-H/SSC-W/SSC-H parameters and the viability dye Aqua Vivid, respectively, and CD4⁺ ($CD3^+CD4^+$) or CD8⁺ ($CD3^+CD4^-$) T-cells were identified based on the CD3/CD4 Abs staining **(A)**. CD4⁺ T-cells were then divided into $FOXP3^+CD127^-$ and $FOXP3^-CD127^+$ CD4⁺ T-cells **(B)**, with Tregs defined as $CD25^+FOXP3^+CD127^-$ **(C).** Further, $CD127^+FOXP3^-$ CD4⁺ T cells were subdivided in central memory (CM) $CCR7^+CD45RA^-$, effector memory (EM) $CCR7^-CD45RA^-$, terminally differentiated EM cells expressing CD45RA (EMRA) $CCR7^-CD45RA^+$, and naïve $CCR7^+CD45RA^+$ cells **(D)**. Shown are histograms reflecting the expression of CCR6, CD26, and CD161 on EM, CM, and EMRA subsets. Th17 cells were identified by Boolean combination of gates between CCR6, CD26 and CD161 and defined as $CCR6^+CD26^+CD161^+$ in each memory subset. Grey histograms represent the expression of CCR6, CD26 and CD161 within the whole population of each memory subset, whereas white histograms illustrate uniquely the positive population for each marker **(E)**.

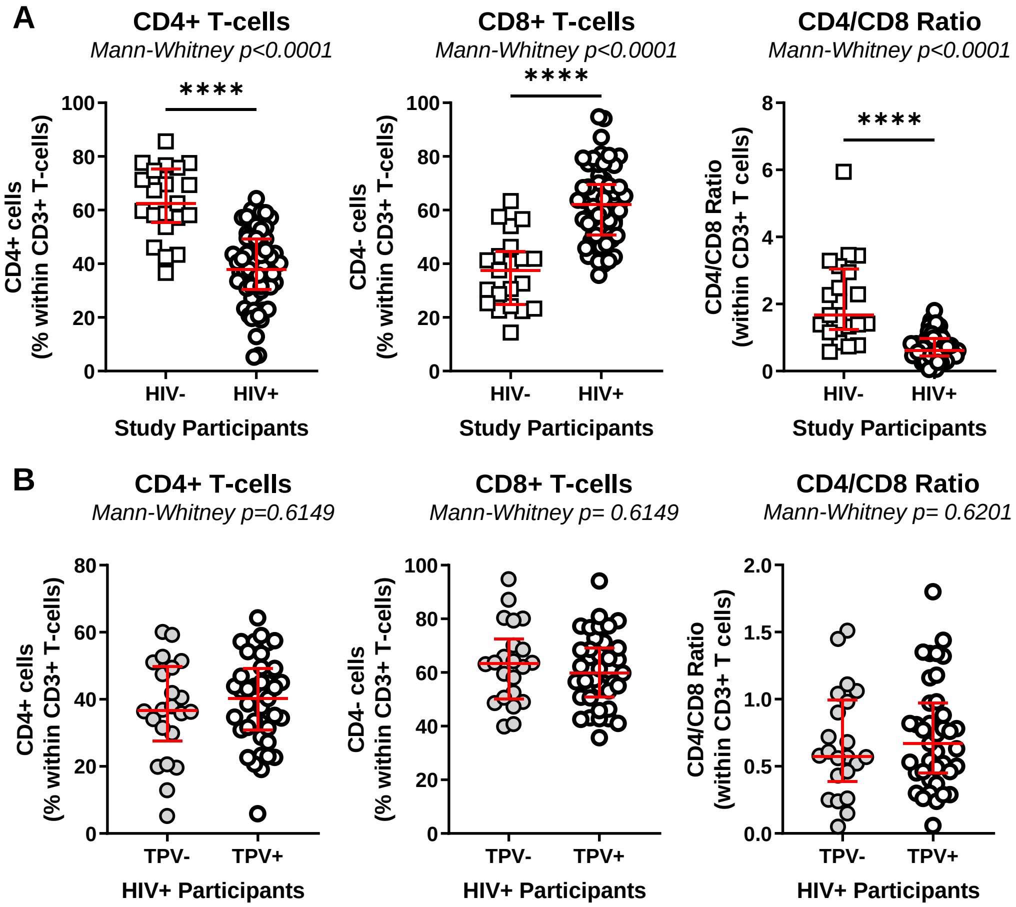


**Figure S4: CD4⁺ and CD8⁺ T-cell frequencies in HIV⁺ and HIV⁻ individuals with/without subclinical atherosclerosis.** T-cell subsets were identified as in **Supplemental Figure 1**. The frequencies of total CD4⁺ (**left panels**) and CD8⁺ T-cells (**middle panels**) and the CD4/CD8 ratios (**right panels**), were compared between HIV⁺ (n=61) and HIV⁻ (n=21) participants **(A)** and among HIV⁺ participants with (TPV⁺; n=39) and without (TPV⁻; n=22) subclinical atherosclerosis **(B)**. Median and IQ range are indicated in red as horizontal lines. Differences among study groups were determined by Mann-Whitney rank test. P-values and statistical significance are indicated in the figures (*, $p < 0.05$; **, $P < 0.01$; ***, $P < 0.001$).

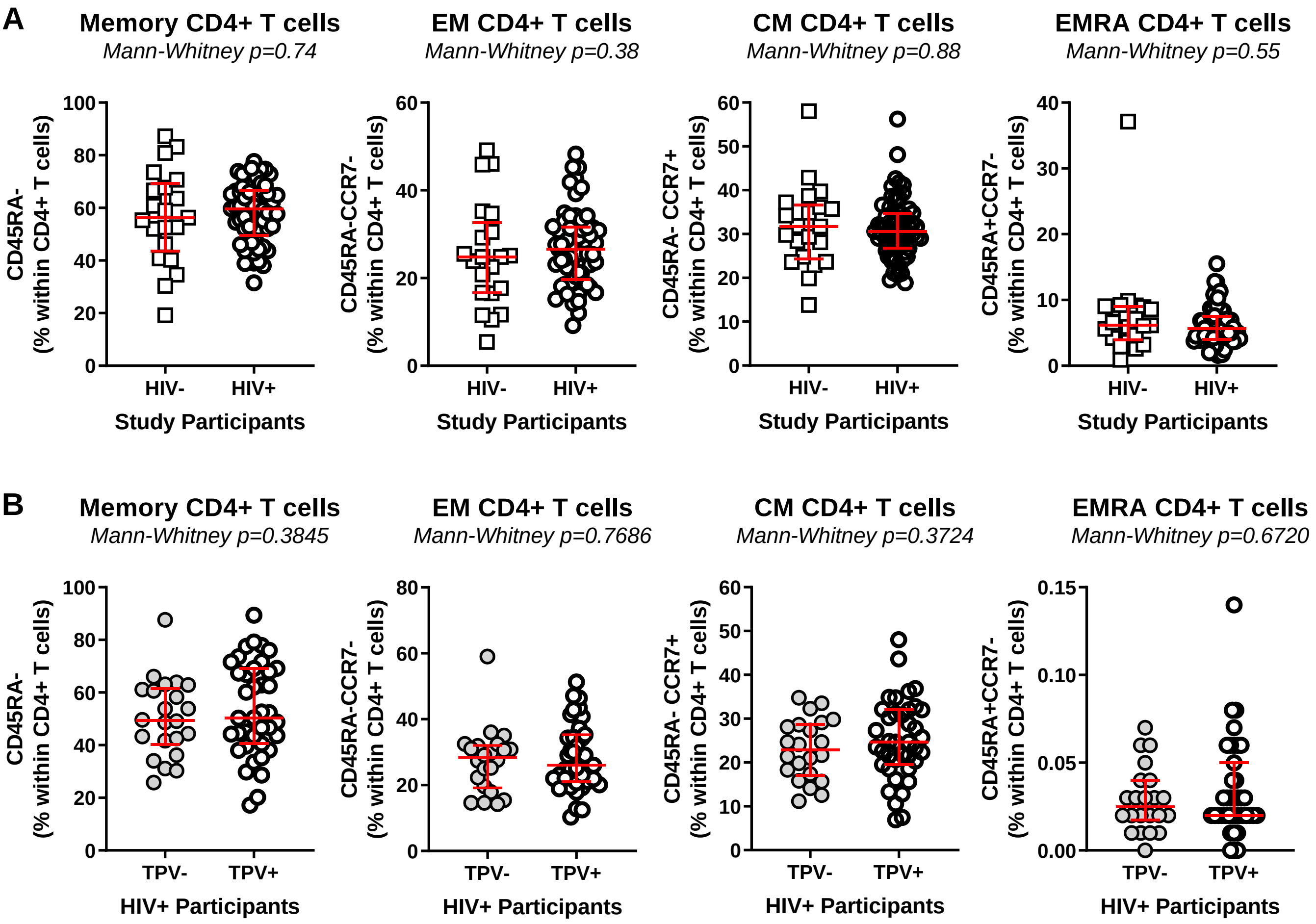


**Figure S5: Frequency of memory CD4$^+$ T-cell subsets in HIV$^-$ and HIV$^+$ participants with and without subclinical signs of atherosclerosis.** T-cell subsets were identified as in **Supplemental Figure 1**. The frequency of CD4$^+$ T cells with a total memory (CD45RA$^-$), as well as EM (CD45RA$^-$CCR7$^-$), CM (CD45RA$^-$CCR7$^+$), EMRA (CD45RA$^+$CCR7$^-$) phenotypes were compared between HIV$^+$ (n=61) and HIV$^-$ (n=21) participants (**A**) and among HIV$^+$ participants with (TPV$^+$; n=39) and without (TPV$^-$; n=22) subclinical atherosclerosis (**B**). Median and IQ range are indicated in red as horizontal lines. Differences among study groups were determined by Mann-Whitney rank test. P-values and statistical significance are indicated in the figures (*, $p < 0.05$; **, $P < 0.01$; ***, $P < 0.001$).

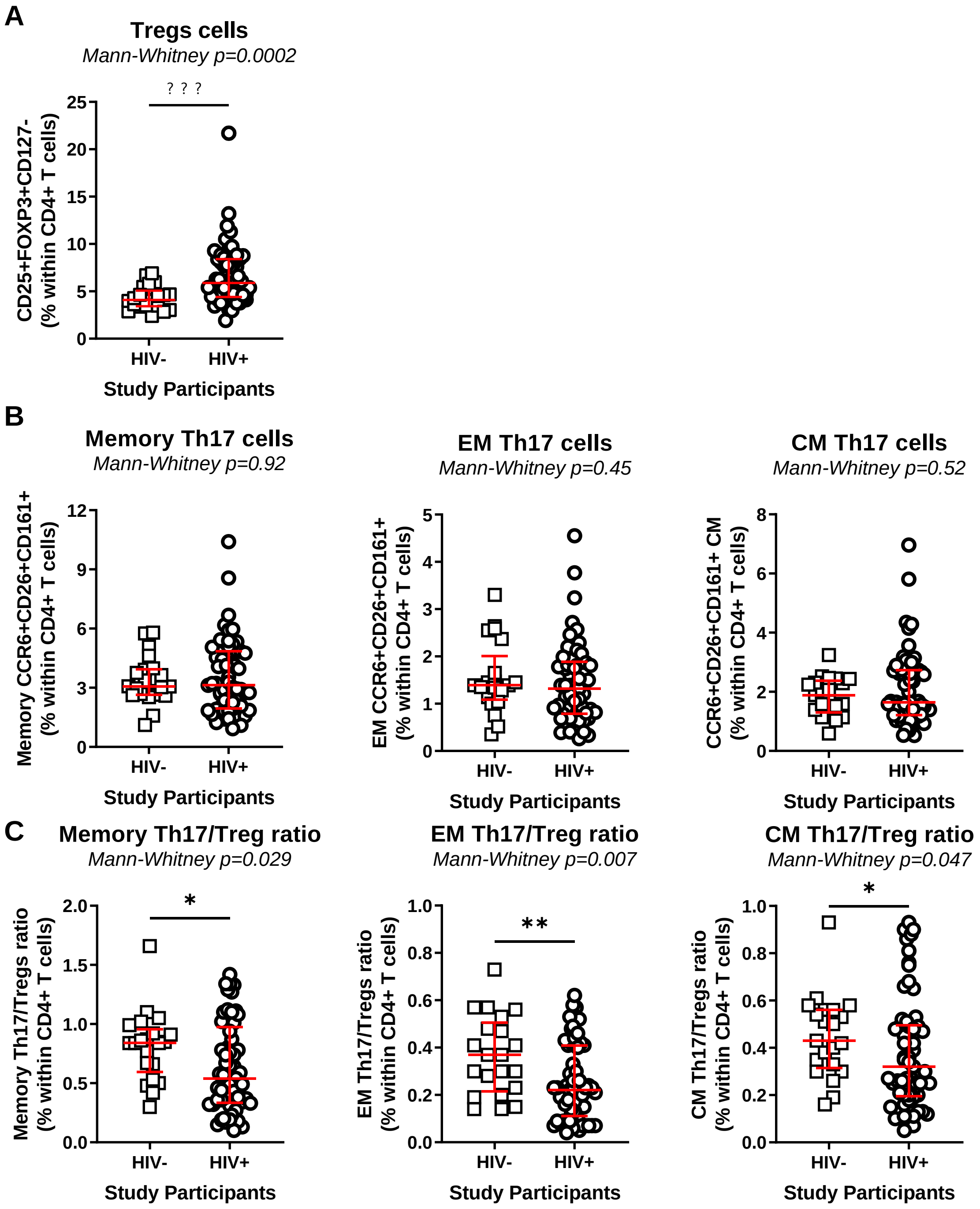


**Figure S6: Frequency of Tregs and Th17 subsets in HIV$^+$ and HIV$^-$ individuals.** The frequencies of Tregs (CD25$^+$FOXP3$^+$CD127$^-$CD4$^+$) **(A)**, as well as Th17-polarized CD4$^+$ T-cells (CCR6$^+$CD26$^+$CD161$^+$) with total memory (CD45RA$^-$), EM (CD45RA$^-$CCR7$^-$) and CM (CD45RA$^-$CCR7$^+$) phenotypes **(B)** were compared between HIV$^+$ and HIV$^-$ participants. **(C)** The Th17/Treg ratios within the memory, EM and CM Th17 subsets were also compared among HIV$^+$ and HIV$^-$ participants. Median values and IQ ranges are indicated in red as horizontal lines. Differences among study groups were determined by Mann-Whitney rank test. P-values and statistical significance is indicated in the figures (*, $p < 0.05$; **, $P < 0.01$; ***, $P < 0.001$). Sample size HIV$^+$ n=61 and HIV$^-$ n=21.

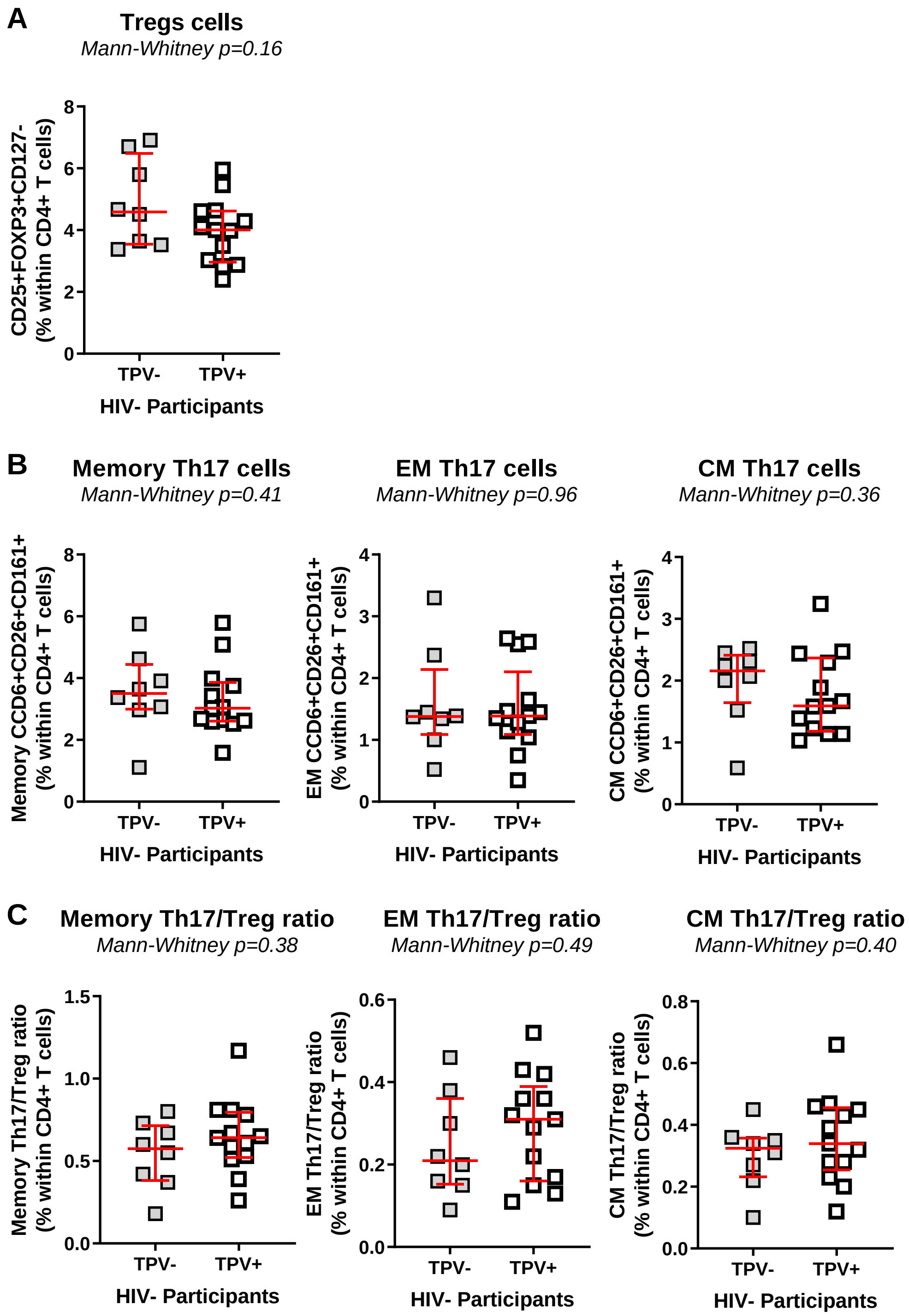


**Figure S7: Frequency of Tregs and Th17 subsets in HIV$^+$ individuals with and without subclinical signs of atherosclerosis.** The frequencies of Tregs (CD25$^+$FOXP3$^+$CD127$^-$CD4$^+$) **(A)**, as well as Th17-polarized CD4$^+$ T-cells (CCR6$^+$CD26$^+$CD161$^+$) with total memory (CD45RA$^-$), EM (CD45RA$^-$CCR7$^-$) and CM (CD45RA$^-$CCR7$^+$) phenotypes **(B)** were compared between HIV$^+$ participants with (TPV$^+$) an without (TPV$^-$) subclinical signs of atherosclerosis. **(C)** The Th17/Treg ratios within the memory, EM and CM Th17 subsets were also compared among TPV$^+$ and TPV$^-$ HIV$^+$ participants. Median values and IQ ranges are indicated in red as horizontal lines. Differences among study groups were determined by Mann-Whitney rank test. P-values and statistical significance is indicated in the figures (*, $p < 0.05$; **, $P < 0.01$; ***, $P < 0.001$). Sample size TPV$^+$ n=39 and TPV$^-$ n=22.

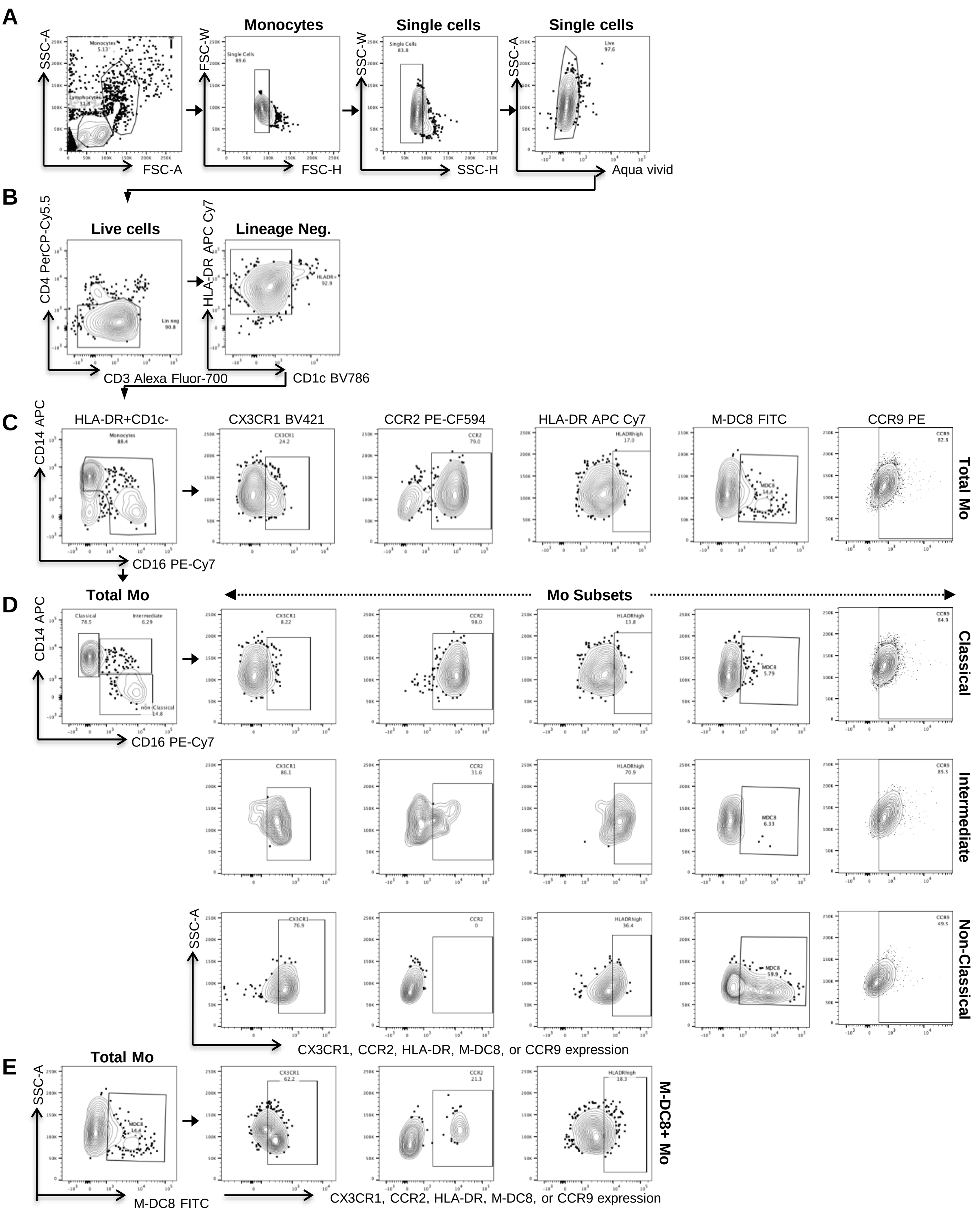


Wiche Salinas et al., Suppl. Figure 8

**Figure S8: Gating strategy for the identification of monocyte subsets**. PBMCs from HIV$^+$ and HIV$^-$ participants were stained with cocktails of fluorochrome-conjugated CD3, CD4, CD1c, CD14, CD16, HLA-DR, CCR6, Slan (M-DC8), CCR2 and CX3CR1 Abs (Monocyte cocktail) or CD3, CD4, CD1C, CD14, CD16, HLA-DR, CCR6, CCR9, CD123 and CD303 Abs (Monocyte and DC cocktail; Supplemental Table 1). Shown is the gating strategy to identify monocyte (Mo) subsets. Briefly, doublets and dead cells were excluded as in Supplemental Figure 3 **(A)**. Total monocytes within cells with a CD4$^-$CD3$^-$CD1c$^-$HLD-DR$^+$ phenotype **(B)** were further divided into classical (CD14$^+$CD16$^-$), intermediate (CD14$^+$CD16$^+$), non-classical (CD14lowCD16$^+$) and Slan/M-DC8$^+$ monocytes **(C-E)**. The expression of CX3CR1, CCR2, CCR9, HLA-DR, and M-DC8 were evaluated on total, classical, intermediate, non-classical and M-DC8$^+$ monocytes. Combination of gates using the Boolean tool for CCR2, CX3CR1, and HLA-DR$^{high}$ or CCR9 and HLA-DR$^{high}$ within monocyte subsets was performed using the single marker gates presented here.

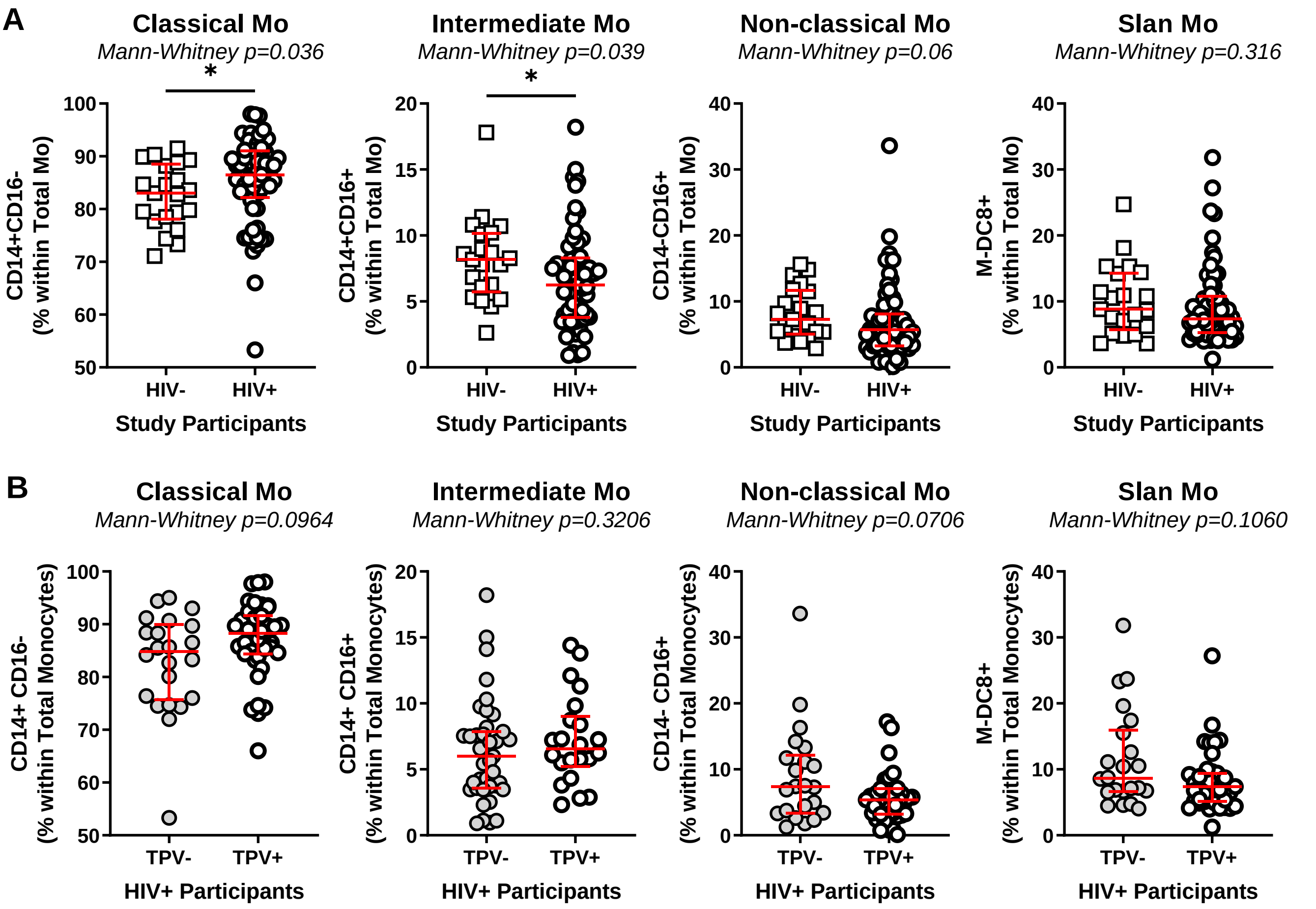


**Figure S9: Frequency of monocyte subsets in HIV$^+$ and HIV$^-$ individuals.**
The frequencies of classical, intermediate, non-classical and Slan/M-DC8+ monocytes were compared between HIV$^+$ (n=61) and HIV$^-$ (n=21) participants **(A),** as well as between TPV$^+$ (n=39) and TPV$^-$ (n=22) HIV$^+$ participants **(B)**. Median and IQ range are indicated in red as horizontal lines. Differences among study groups were determined by Mann-Whitney rank test. P-values and statistical significance is indicated in the figures (*, $p < 0.05$; **, $P < 0.01$; ***, $P < 0.001$).

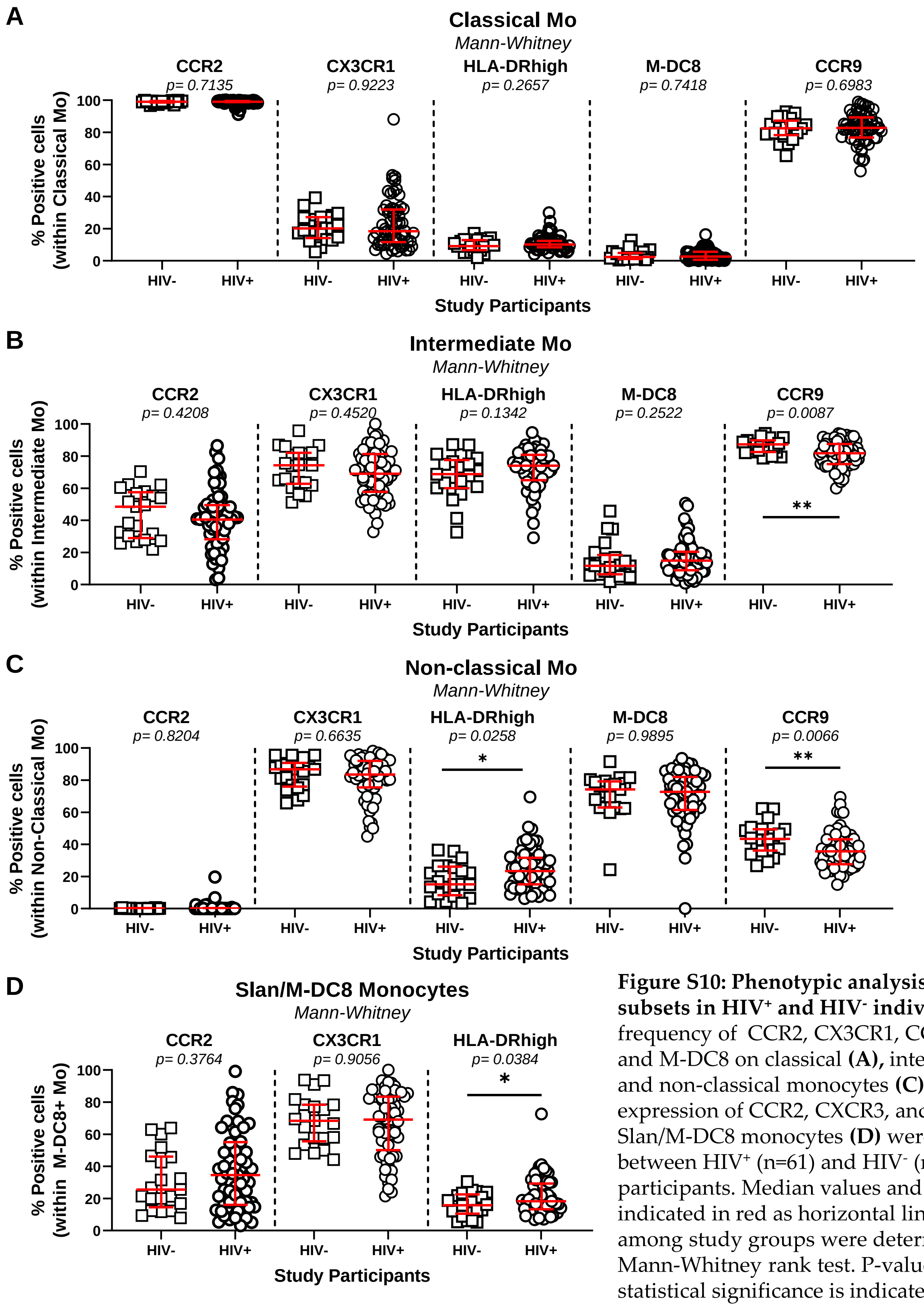


**Figure S10: Phenotypic analysis of monocyte subsets in HIV+ and HIV- individuals.** The frequency of CCR2, CX3CR1, CCR9, HLA-DR, and M-DC8 on classical **(A),** intermediate **(B)**, and non-classical monocytes **(C)**, as well as the expression of CCR2, CXCR3, and HLA-DR on Slan/M-DC8 monocytes **(D)** were compared between HIV+ (n=61) and HIV- (n=21) participants. Median values and IQ ranges are indicated in red as horizontal lines. Differences among study groups were determined by Mann-Whitney rank test. P-values and statistical significance is indicated in the figures (*, p < 0.05; **, P < 0.01; ***, P < 0.001).

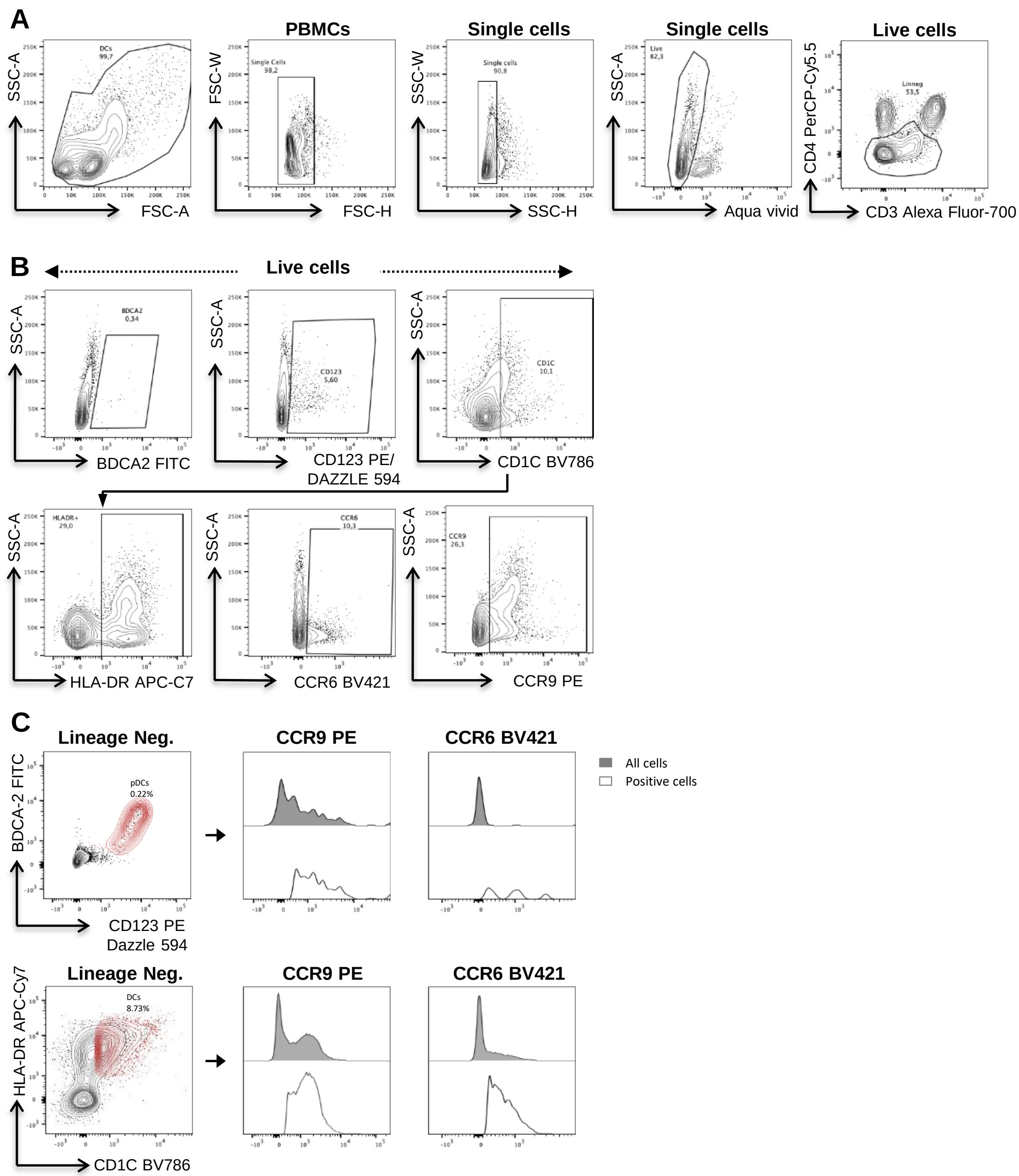


**Figure S11. Gating strategy for the identification of DC subsets.** PBMCs from HIV$^+$ and HIV$^-$ participants were stained with a cocktail of fluorochrome-conjugated CD3, CD4, CD1C, CD14, CD16, HLA-DR, CCR6, CCR9, CD123 and BDCA2 Abs (Supplemental Table 1). Show is the gating strategy to identify plasmacytoid (pDCs) and myeloid DC (mDC). **(A)** Doublets and dead cells were excluded, and Lineage negative (Lin$^{neg}$) cells were determined as CD3$^{low/-}$ and CD4$^{low/-}$. **(B)** Shown are gates to determine the surface expression of BDCA2, CD123, CD1C, HLA-DR, and CCR6 within Live cells. **(C)** Boolean gating using gates set in B were used to identify pDCs (CD123$^+$CD303$^+$Lin$^{neg}$) (top panels) and mDCs (HLA-DR$^+$CD1C$^+$Lin$^{neg}$) (bottom panels). CCR6 and CCR9 expression was evaluated in both subsets. Grey histograms represent the expression of CCR6 and CCR9 within the whole population of each memory subset, whereas white histograms illustrate uniquely the positive population for each marker.

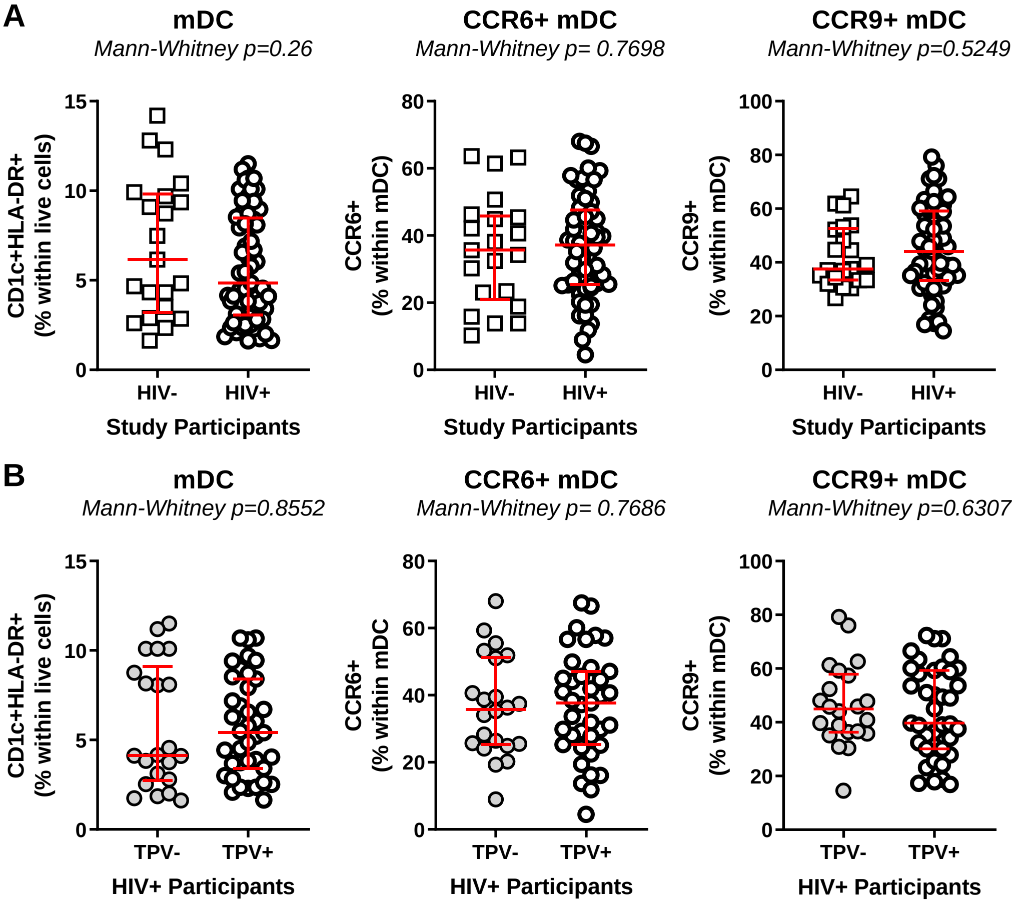


**Figure S12. Dynamics of mDC in HIV⁻ and HIV⁺ individuals with or without coronary plaque.** The frequencies of total, CCR6$^{+}$ and CCR9$^{+}$ mDC (CD11c$^{+}$HLA-DR$^{+}$) were compared between HIV$^{+}$ (n=61) and HIV$^{-}$ (n=21) participants **(A)**, as well as between TPV$^{+}$ (n=39) and TPV$^{-}$ (n=22) HIV$^{+}$ participants **(B)**. Median values and IQ ranges are indicated in red as horizontal lines. Differences among study groups were determined by Mann-Whitney rank test. P-values and statistical significance are indicated in the figures (*, $p < 0.05$; **, $P < 0.01$; ***, $P < 0.001$).

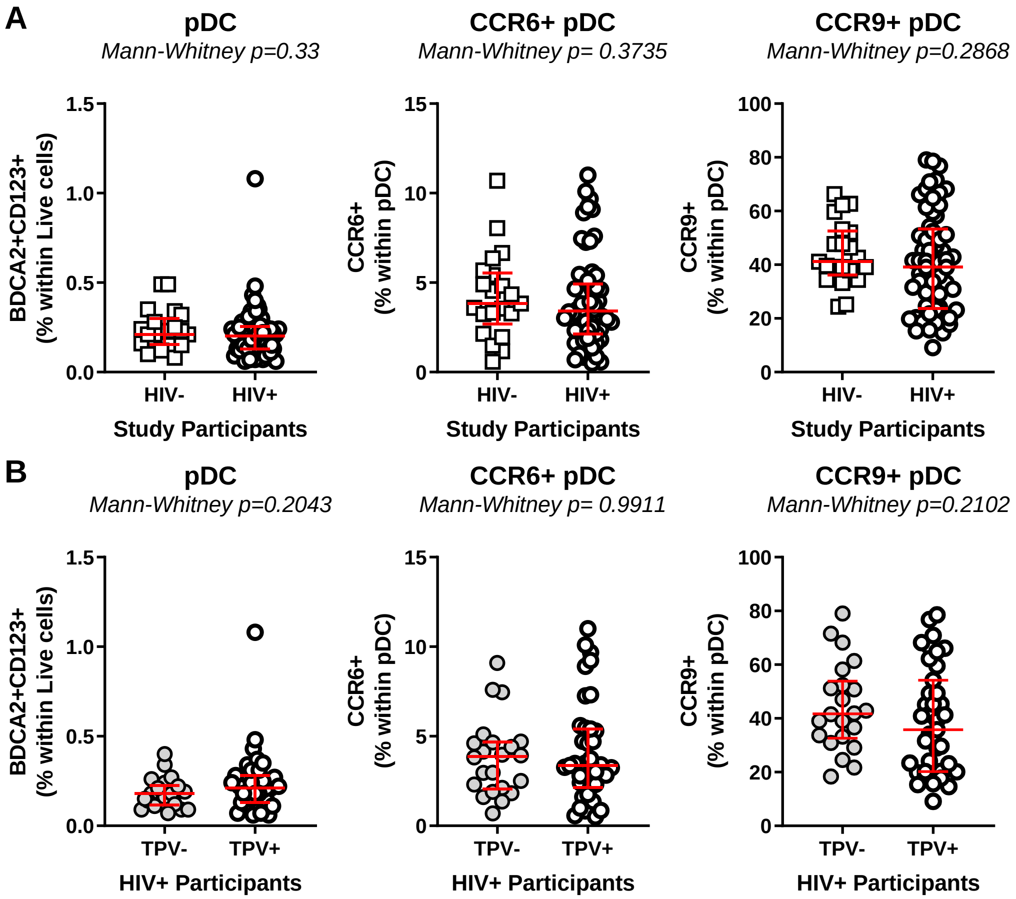


**Figure S13. Dynamics of pDC in HIV$^-$ and HIV$^+$ individuals with or without coronary plaque.** The frequencies of total, CCR6$^+$ and CCR9$^+$ pDC (BDCA2$^+$CD123$^+$) were compared between HIV$^+$ (n=61) and HIV$^-$ (n=21) participants **(A)**, as well as between TPV$^+$ (n=39) and TPV$^-$ (n=22) HIV$^+$ participants **(B)**. Median values and IQ ranges are indicated in red as horizontal lines. Differences among study groups were determined by Mann-Whitney rank test. P-values and statistical significance are indicated in the figures (*, $p < 0.05$; **, $P < 0.01$; ***, $P < 0.001$).
